# Microglia transcriptomic response to doxorubicin-induced genotoxic stress

**DOI:** 10.64898/2026.10.05.756646

**Authors:** Ekkaphot Khongkla

## Abstract

Chemobrain, characterized by cognitive dysfunction, has emerged as a significant health concern following cancer treatment. Doxorubicin, an FDA-approved chemotherapeutic agent, is widely used to treat various cancers. However, its adverse effects raise concerns regarding toxicity in non-cancerous cell types. Microglia, the resident immune cells of the brain, play a central role in maintaining neural homeostasis. Despite this, the mechanisms by which microglia become dysfunctional following doxorubicin exposure remain poorly understood. Understanding the transcriptomic alterations induced by doxorubicin in microglia could provide critical insights into mechanisms underlying its neurotoxic effects. In this study, the transcriptomic response of human microglia to doxorubicin using mRNA sequencing combined with bioinformatic analysis was investigated. Exposure to doxorubicin significantly altered microglial morphology, promoting a shift toward a process-bearing phenotype in human HMC3 cells. This phenotypic change was accompanied by elevated levels of DNA damage and was associated with the induction of cellular senescence. At the gene regulation level, the Illumina-based transcriptomic analysis provided 25,834 gene expression profiling in which 5,669 were differentially expressed. The future analyses revealed that 3,431 (upregulated) and 2,238 (downregulated) were differentially expressed under the treatment of doxorubicin. Genes associated with the p53 signaling pathway and inflammatory responses were upregulated, whereas genes involved in translation, DNA replication, and cell-cycle progression were downregulated. This transcriptional dysregulation was accompanied by differential expression of key transcription factors. This study provides transcriptomic evidence elucidating the genotoxic response to doxorubicin in microglial cells, offering mechanistic insights into chemobrain and highlighting key molecular pathways as potential targets for therapeutic intervention.

**Highlights:**

- Doxorubicin induces persistent DNA damage and senescence-associated phenotypic changes in human HMC3 microglial cells.
- Unbiased Illumina-based transcriptomic profiling identified 25,834 expressed genes, including 5,669 differentially expressed genes following doxorubicin treatment.
- Genes involved in p53 signaling and inflammatory responses were upregulated, whereas genes associated with translation, DNA replication, and cell -cycle progression were downregulated.
- Transcriptomic dysregulation was accompanied by altered expression of key transcription factors, including interferon regulatory factor 4 (IRF4).
- These findings provide a transcriptomic framework for investigating the potential contribution of microglial dysfunction to chemotherapy-associated neurotoxicity.

## 1. Introduction

Chemotherapy-related cognitive impairment, commonly referred to chemobrain, is a prevalent and clinically significant complication among cancer survivors following chemotherapy. Cognitive impairment has been reported in up to 75% of patients shortly after treatment(Neth et al., 2026). Affected patients may experience a range of cognitive deficits, including impairments in memory, attention, executive function, and learning ability, which can substantially affect daily functioning and quality of life. Importantly, these cognitive effects may occur independently of direct tumor-related processes, suggesting that chemotherapy-induced dysfunction of non-malignant brain cells may contribute to the development and progression of cognitive impairment.

Doxorubicin is an anthracycline chemotherapeutic agent widely used in the treatment of breast cancer, bladder cancer, lymphoma and acute lymphocytic leukemia . Its antineoplastic activity is mediated through multiple mechanisms including DNA intercalation, inhibition of topoisomerase II, and generation of reactive oxygen species, collectively resulting in DNA damage and inhibition of cancer cell proliferation(Baxter-Holland and Dass, 2018; C. Gao et al., 2023). These biological effects are not exclusively restricted to cancer cells, and exposure to doxorubicin can also induce genotoxic stress in healthy tissues. The ability of doxorubicin to affect the CNS has raised concerns regarding its potential contribution to chemotherapy-associated neurological complications. The underlying mechanisms studied so far usually focus on neurons (Kamińska and Cudnoch-Jędrzejewska, 2023), while the cellular and molecular responses of other brain-resident cells to doxorubicin remain incompletely characterized.

Microglia are the resident macrophages of the CNS, comprising approximately 10–15% of all glial cells, and play essential roles in immune surveillance, synaptic homeostasis, tissue repair, and maintenance of the neural microenvironment(Carr et al., 2025; C. Gao et al., 2023). In response to cellular stress or injury, microglia undergo dynamic morphological and functional changes accompanied by extensive molecular reprogramming. Dysregulated microglial activation can contribute to neuronal dysfunction and neuroinflammation. Emerging evidence suggests that chemotherapeutic agents, including doxorubicin, may affect microglial function(Barbosa-Azevedo et al., 2024; Demby et al., 2024; Marques et al., 2020). However, the molecular mechanisms underlying doxorubicin-induced microglial dysfunction remain poorly understood. In particular, a comprehensive characterization of the transcriptional alterations induced by doxorubicin in human microglial cells is lacking. Therefore, systematic analysis of the microglial transcriptome following doxorubicin exposure is necessary to identify molecular pathways and regulatory mechanisms that may contribute to chemotherapy-associated neurotoxicity.

mRNA sequencing (mRNA-seq) has emerged as a powerful approach for unbiased, genome-wide profiling of gene expression(Lowe et al., 2017). Unlike targeted analyses, RNA-seq enables comprehensive characterization of transcriptional alterations across thousands of genes and facilitates the identification of differentially expressed genes, enriched biological pathways, and transcriptional regulatory networks in response to specific cellular conditions(Hassan, 2023; Ibanez et al., 2026; Noorbakhsh et al., 2015). Integration of RNA-seq with bioinformatic analyses, including pathway enrichment and transcription factor analysis, can therefore provide mechanistic insights into cellular responses to pharmacological stress and reveal molecular processes that may not be apparent from individual candidate-gene studies(Anders and Huber, 2010; Kanehisa and Goto, 2000; Young et al., 2010).

Therefore, this study aimed to characterize the transcriptomic response of human microglial cells to doxorubicin exposure using mRNA sequencing coupled with bioinformatic analyses. Human HMC3 microglial cells were exposed to doxorubicin, followed by assessment of cellular morphological changes, DNA damage, and cellular senescence. To determine key transcriptional regulators associated with the cellular response to doxorubicin, bulk transcriptomic profiling was subsequently performed to identify differentially expressed genes and dysregulated biological pathways. These findings provide molecular insights into doxorubicin-induced microglial dysfunction and may contribute to a better understanding of the cellular and molecular mechanisms underlying chemobrain.

## 2. Materials and methods

### 2.1 Reagent

Doxorubicin (cat.no. 5927, Cell Signaling Technology, USA) was prepared at 10 mM in DMSO. The stock was diluted into 1 mM as a working stock. Antibodies used in this study is listed in Supplementary Table 1. Human primer oligonucleotide sequences are listed in Supplementary Table 2.

### 2.2 Cell culture

Human microglial cells (HMC3) were obtained from American Type Culture Collection (ATCC, CRL-3304). The cells were cultured under Minimum Essential Medium Eagle (MEM, Cat.no.10-010-CV, Corning, USA) supplemented with 10 % fetal bovine serum (Cat.no. 35-078-CV, Corning) and 1 % Penicillin-streptomycin (10,000 U/mL) (cat.no.15140122, Gibco, USA). Cells were maintained in a humidified incubator with 5% CO_2_ at 37 °C.

### 2.3 Cell viability measurement

Cells at the density of 10^4^ cells/well in a 96-well plate were exposed to different concentrations of the doxorubicin (0.01-5 μM) in complete media. In the protocol 1, cell viability was measured after 24 h of drug treatment. In Protocol 2, the drug-containing media were removed and replaced with the fresh complete media for 48 h before measuring cell viability. MTT assay (M5655, Sigma‒Aldrich, USA) was used to measure cell viability. The MTT stock reagent (5 mg/ml) was added to the conditioning media to obtain the final MTT reagent concentration (0.5 mg/ml). The plates were further cultured for additional 3 h. The MTT solution was removed, and insoluble formazan was dissolved in DMSO. Colorimetric determination was performed at 570 nm via an EZ Read2000 microplate reader (Biochrom, USA). The data were normalized to the control data.

### 2.4 Immunoblotting

HMC3 cells were seeded at the density of 10^6^ cells at a 75 cm^2^ culture dish. The plated cells were exposed to DMSO (vehicle control) or Doxorubicin (1 μM) for 24 h. Cell pellets were collected and total proteins were isolated with RIPA lysis buffer containing 10% Triton X-100. Soluble fraction was obtained after centrifugation at 15,000 *xg*. The concentration of the extracted proteins was measured via a Pierce BCA protein assay kit (cat no. 23225, Thermo Scientific, USA). An identical concentration of total extracted protein was prepared in 1X Laemmli SDS sample buffer (Thermo Scientific, USA), heated at 95°C for 15 min, separated by SDS‒PAGE, and transferred to a PVDF membrane (1620177, Bio-Rad Laboratories, USA). The non-specific proteins were blocked with 2.5% nonfat dry milk in TBST for 30 min and incubated with primary antibodies against, SIRT1, γH2AX or GAPDH (Supplementary Table1). HRP-conjugated secondary antibodies for anti-rabbit (7074, Cell Signaling, USA) IgG were used for detection with Pierce enhanced chemiluminescence (ECL) western blot substrate (cat.no. 32209, Thermo Scientific, USA). The signals were imaged with a Fusion FX chemiluminescence system (Vilber, France). Uncropped western blot images are provided in Supplementary File 7.

### 2.5 Immunocytochemistry

The cells were fixed with 4% paraformaldehyde (PFA/ 1XPBS) for 15 min at room temperature and rinsed three times with PBS. The cells were then permeabilized in 0.2% Triton X-100 at room temperature. The plates were washed with PBS three times. Nonspecific binding was blocked with 1% bovine serum albumin (BSA) in PBS for 30 min at 37°C. The cells were stained with the primary antibody against Iba1, γH2AX or Ki67 (Supplementary Table 1) on parafilm for 30 min at 37°C before being washed with PBS three times, each for more than 5 min. The goat anti-rabbit IgG (H+L) cross-adsorbed secondary antibody, Alexa Fluor 488 (A32790, Invitrogen, USA), was incubated for 30 min at 37°C. The incubated cells were washed with PBS three times, dipped in clean water, and mounted on slides using Fluoromount-G Mounting Medium with DAPI (cat no. 00495952; Invitrogen, USA). Images were acquired via a Zeiss LSM 800 confocal laser scanning microscope.

### 2.6 Senescence-associated β-galactosidase detection

The senescent cells were detected using Senescence β-galactosidase staining kit (cat. no. 9860, Cell Signaling Technology, USA). The culture media were removed and the cells were washed one time with 1XPBS. The 1X fixative solution was added to fix the cells for 15 min at RT. The cells were washed twice with 1X PBS. The β-galactosidase staining solution containing 50 μL of 20 mg/mL X-gal stock solution was added to the cells. to prevent evaporation, the plate was sealed with parafilm and incubated in a dry incubator at 37 °C overnight. The presence of β-galactosidase -positive cells was observed under the light microscope.

### 2.7 Quantitative real-time polymerase chain reaction (qRT‒PCR)

Total RNAs were extracted from the microglial HMC3 cells using PureLink RNA mini kit (cat.no. 12183018A, Invitrogen, USA) according to the manufacturer’s instructions. RNA quantitation was performed by NanoDrop One (Thermo Fisher Scientific, USA). cDNA synthesis was conducted via an iScript cDNA Synthesis Kit (Bio-Rad Laboratories, USA). Total cDNA (100 ng in 2 µl) was mixed with 8 µl of iTaq Universal SYBR Green Supermix (Bio-Rad Laboratories, USA) containing specific forward and reverse primers (400 nM each) to perform real-time PCR amplification, following the manufacturer’s instructions. The relative mRNA expression of the target genes was normalized to that of GAPDH, which served as the internal control and was calculated via the 2 ^-ΔΔCt^ method. A list of primers used in this study is provided in Supplementary Table 2.

### 2.8 mRNA-sequencing analysis

The procedure for total RNA extraction follows the previous section. RNA integrity was assessed using the gel agarose electrophoresis (Supplementary Fig. 1), and the RNA Nano 6000 Assay Kit of the Agilent Bioanalyzer 2100 system (Agilent Technologies, USA).

#### 2.8.1 Library preparation for Transcriptome sequencing

A total amount of 1 μg RNA per sample was used as input material for the RNA sample preparations. Sequencing libraries were generated using NEBNext UltraTM RNA Library Prep Kit for Illumina (NEB, USA) following manufacturer’s recommendations and index codes were added to attribute sequences to each sample. Briefly, mRNA was purified from total RNA using poly-T oligo-attached magnetic beads. Fragmentation was carried out using divalent cations under elevated temperature in NEBNext First Strand Synthesis Reaction Buffer (5X). First strand cDNA was synthesized using random hexamer primer and M-MuLV Reverse Transcriptase. Second strand cDNA synthesis was subsequently performed using DNA Polymerase I and RNase H. Remaining overhangs were converted into blunt ends via exonuclease/polymerase activities. After adenylation of 3’ ends of DNA fragments, NEBNext Adaptor with hairpin loop structure were ligated to prepare for hybridization. In order to select cDNA fragments of preferentially 240 bp in length, the library fragments were purified with AMPure XP system (Beckman Coulter, Beverly, USA). Then 3 μl USER Enzyme (NEB, USA) was used with size-selected, adaptor-ligated cDNA at 37°C for 15 min followed by 5 min at 95°C before PCR. Then, PCR was performed with Phusion High-Fidelity DNA polymerase, Universal PCR primers and Index (X) Primer. At last, PCR products were purified (AMPure XP system) and library quality was assessed on the Agilent Bioanalyzer 2100 system. The clustering of the index-coded samples was performed on a cBot Cluster Generation System using TruSeq PE Cluster Kit v4-cBot-HS (Illumia) according to the manufacturer’s instructions. After cluster generation, the library preparations were sequenced on an Illumina platform and paired-end reads were generated.

#### 2.8.2 Data analysis

Raw data (raw reads) of fastq format were firstly processed through in-house perl scripts. In this step, clean data (clean reads) were obtained by removing reads containing adapter, reads containing ploy-N and low-quality reads from raw data. At the same time, Q20, Q30, GC-content and sequence duplication level of the clean data were calculated. All the downstream analyses were based on clean data with high quality. The adaptor sequences and low-quality sequence reads were removed from the data sets. Raw sequences were transformed into clean reads after data processing. These clean reads were then mapped to the reference genome sequence. Only reads with a perfect match or one mismatch were further analyzed and annotated based on the reference genome. Hisat2 tools soft were used to map with reference genome. Gene function was annotated based on the databases indicated in Supplementary Table 6. Gene expression levels were estimated by fragments per kilobase of transcript per million fragments mapped. The formula is shown as follow:

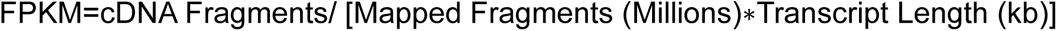

#### 2.8.3 Differential expression analysis

Differential expression analysis of two groups was performed using the DESeq2. DESeq2 provide statistical routines for determining differential expression in digital gene expression data using a model based on the negative binomial distribution. The resulting P values were adjusted using the Benjamini and Hochberg’s approach for controlling the false discovery rate. Genes with an adjusted P-value < 0.01 found by DESeq2 were assigned as differentially expressed. Gene Ontology (GO) enrichment analysis of the differentially expressed genes (DEGs) was implemented by the GOseq R packages based Wallenius non-central hyper-geometric distribution (Young et al., 2010), which can adjust for gene length bias in DEGs. Transcription Factor (TFs) of DEGs were identified and annotation based on AminalTFDB database. Transcription factor binding site (TFBS) in promoter regions of DEGs was predicted by R package TFBSTools (Tan and Lenhard, 2016). JASPAR database was used as reference motif database (Fornes et al., 2020).

### 2.9 Statistics

GraphPad Prism version 8 (GraphPad Software, USA) was used for statistical analyses. Differences between two groups were analyzed using an unpaired Student’s t-test, while comparisons among multiple groups were performed using one-way analysis of variance (ANOVA). A p-value < 0.05 was considered statistically significant. The number of biological replicates is specified in the corresponding figure legends.

## 3. Result

### 3.1 Doxorubicin induces irreversible loss of cell viability and morphological alterations in HMC3 cells

To investigate the cytotoxic effects of doxorubicin, HMC3 cells were exposed to various concentrations (0.01, 0.05, 0.1, 0.5, 1, and 5 μM) according to Protocols #1 and #2 (Fig. 1A). The results showed that doxorubicin at 1 and 5 μM reduced cell viability by approximately 20% (Fig. 1B). More pronounced effects were observed in Protocol #2 (Fig. 1C), suggesting that the growth arrest could not be reversed after replacement with fresh medium. Since both concentrations (1 and 5 μM) produced similar effects, the lower concentration (1 μM) was selected for subsequent experiments.

**Figure 1.**
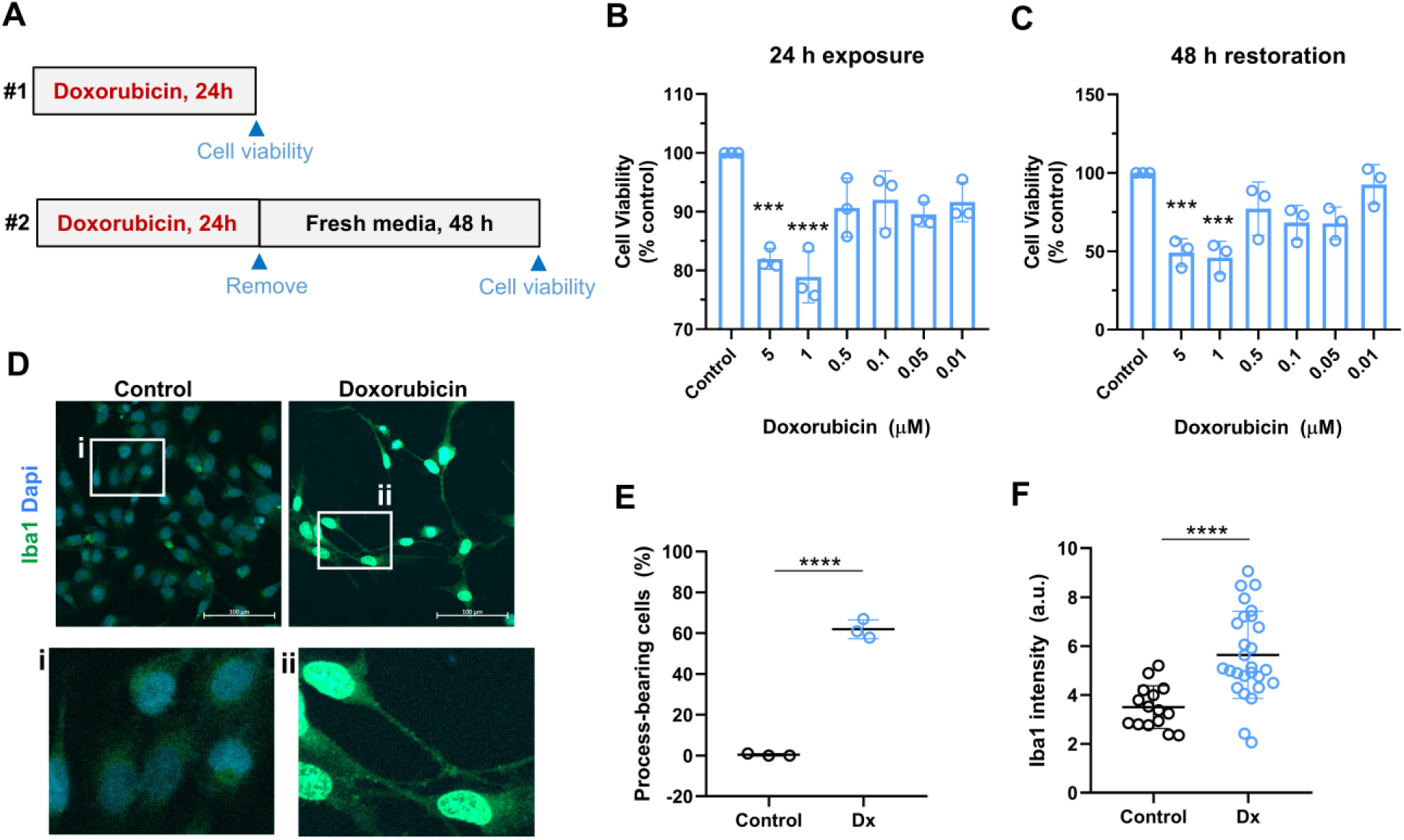
Doxorubicin reduces cell viability and promotes morphological changes toward a process-bearing phenotype in human microglial HMC3 cells. **(A)** Schematic illustration of the two doxorubicin exposure protocols (#1 and #2) used to investigate the effects of doxorubicin at different concentrations (0.01–5 μM) on the viability of human HMC3 cells. Cell viability was quantitatively assessed using the MTT assay following protocol #1 **(B)** and protocol #2 **(C)** and expressed relative to the untreated control. Data are presented as percentages normalized to the control and shown as means ± SD from three biological replicates (n = 3). Statistical significance among experimental groups was determined using one-way ANOVA. ***p < 0.001 and ****p < 0.0001 versus control. **(D)** Representative immunofluorescence images of HMC3 cells stained for Iba1 (green) and DAPI (blue) under unstimulated and doxorubicin-treated conditions (1 μM). Enlarged images (i and ii) correspond to the highlighted regions. Scale bars = 100 μm. **(E)** Quantification of process-bearing HMC3 cells in doxorubicin-treated (Dx) groups relative to the control (%). **(F)** Quantification of Iba1 fluorescence intensity in doxorubicin-treated and control groups. Data are presented as means ± SD, with each dot representing an independent biological replicate. Statistical significance between the two groups was determined using an unpaired Student’s t-test. ****p < 0.0001.

Under doxorubicin treatment conditions, noticeable morphological changes in the microglia cells were observed. To further characterize these alterations, immunocytochemical analysis of Iba1, a well-established microglial marker, was performed. Doxorubicin-treated cells exhibited a more elongated morphology with prominent processes compared to untreated controls (Fig. 1D–E). In addition, Iba1 intensity was significantly increased following treatment (Fig. 1F). Collectively, these results demonstrate that doxorubicin induces a clear morphological response in human microglial HMC3 cells.

### 3.2 Doxorubicin triggers DNA damage and senescent characteristics in HMC3 cells

Doxorubicin, a potent anthracycline chemotherapeutic agent, exerts its effects by intercalating into DNA and inhibiting topoisomerase II, thereby inducing genotoxic stress in various cell types (Y. Gao et al., 2023; Kamińska and Cudnoch-Jędrzejewska, 2023; Marques et al., 2020; Patel et al., 2025). However, whether this drug promotes DNA damage in human microglia remains unclear. Immunocytochemical staining for nuclear γH2AX, a marker of DNA damage, revealed an increased γH2AX intensity in doxorubicin-treated cells (Fig. 2A-B). Sirtuin-1 (SIRT1) is a NAD+-dependent deacetylase crucial for genome stability, acting as a key regulator in the DNA damage response and repair. As shown in Figure 2C-E, doxorubicin treatment significantly increased SIRT1 expression, concomitant with elevated γH2AX levels. This suggests that microglial cells may attempt to mitigate DNA damage by upregulating a SIRT1 - mediated adaptive response.

**Figure 2.**
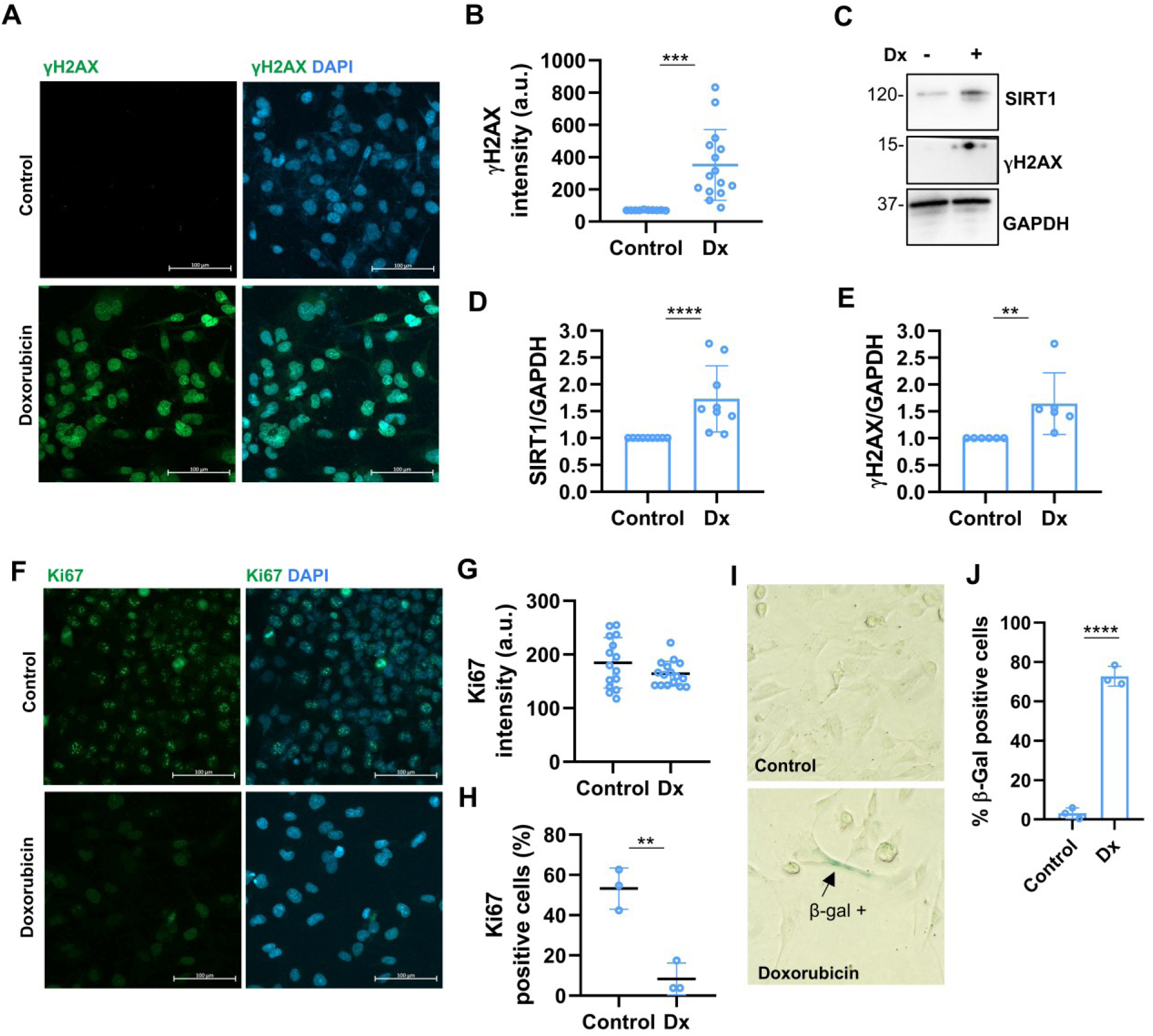
Doxorubicin induces genotoxic stress response accompanied by DNA damage, SIRT1 upregulation, and senescence-associated phenotypes in human HMC3 cells. **(A)** Representative immunofluorescence images of HMC3 cells stained for γH2AX, a marker of DNA damage (green) and DAPI (blue) under unstimulated and doxorubicin (Dx)-treated conditions (1 μM). Scale bar is 100 μm. **(B)** Intensity of γH2AX quantified from the immunocytochemistry (% control). **(C)** Representative western blot images showing the increased level of SIRT1 and γH2AX in the Dx -treated cells. Quantification of SIRT1 **(D)** and γH2AX **(E)** from the western blot analysis. Each dot represents different biological replicate (n=6-9). **(F)** Representative immunofluorescence images of HMC3 cells stained for ki67, a proliferative marker (green) and DAPI (blue) under the unstimulated and Dx-treated conditions. Scale bar is 100 μm. **(G)** Intensity of Ki67, and **(H)** a number of ki67-positive cells (% of total cells) quantified from the immunocytochemistry. **(I)** Representative images of β-galactosidase positive cells and **(J)** their quantitation normalized to total cells (n=3). Statistical significance between the two groups was determined using an unpaired Student’s t-test. **p< 0.01, ***p<0.001, ****p < 0.0001.

DNA damage is a primary driver of cellular senescence, a state of cell cycle arrest that contributes to age-related diseases. As an indicator of cellular senescence, nuclear Ki67 which is a marker of active cell proliferation and reduced in senescent cells, was evaluated. The data show that doxorubicin significantly decreased the proportion of Ki67-positive cells, confirming cell cycle arrest following doxorubicin-induced genotoxic stress in HMC3 cells (Fig. 2F–H). The induction of senescence by doxorubicin was further corroborated by an increased number of β-galactosidase–positive cells (Fig. 2I–J). Collectively, these findings demonstrate that doxorubicin induces pronounced DNA damage in human microglial HMC3 cells and promotes the development of a senescence-associated phenotype.

### 3.3 mRNA-seq reveals extensive gene expression changes following doxorubicin treatment in HMC3 cells

While the phenotypic effects of doxorubicin on microglial cells suggest activation of DNA damage and senescence pathways, the transcriptional regulatory profile underlying these processes remains unclear. To investigate the microglial response to doxorubicin at the gene expression level, transcriptomic analysis was performed to obtain an unbiased and holistic profile of global gene expression (Fig.3A). The cells were treated with doxorubicin in three biological replicates following the described protocol. Total RNA was then extracted, and its integrity was evaluated by gel electrophoresis (Supplementary Fig. 1). The presence of two intense rRNA bands (28S and 18S) indicated high RNA integrity, confirming that the samples were suitable for mRNA-seq analysis.

**Figure 3.**
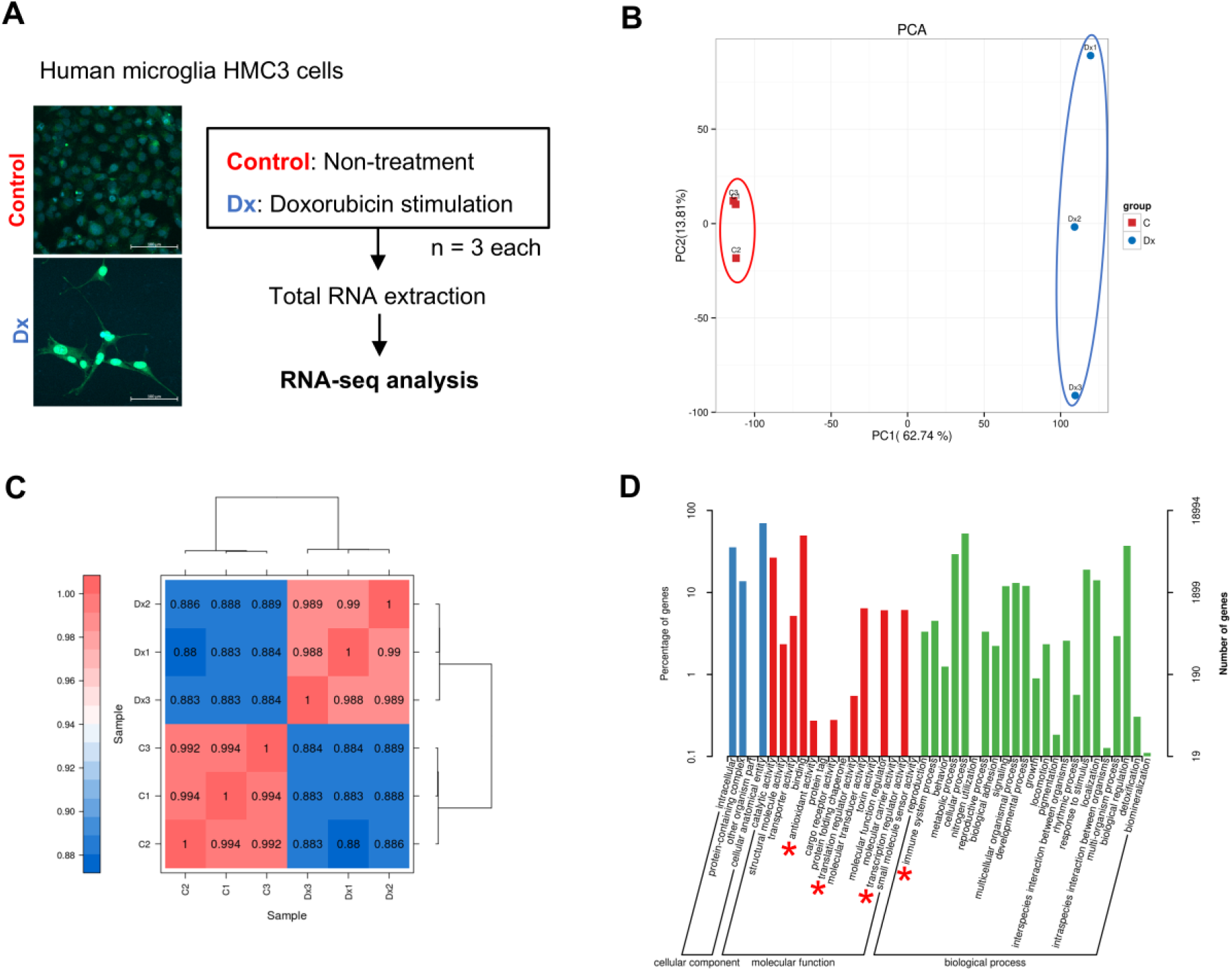
Doxorubicin induces substantial change of gene expression in human microglial HMC3 cells. **(A)** Illustration summarizes the mRNA -seq workflow. **(B)** Principal component analysis (PCA) and **(C)** sample correlation of transcriptomic profiles between control (c) and Doxorubicin -treated cells (Dx). Each dot represents each biological replicate (n=3). The first two principal components (PC1 and PC2) account for 62.74% and 13.81% of the variance, respectively, indicating clear separation among groups. Heatmap of sample‒ to‒sample Pearson correlation coefficients with hierarchical clustering. High intragroup similarity and distinct intergroup clustering confirm the reliability and reproducibility of the RNA sequencing data. **(D)** Gene Ontology classification of identified genes according to cellular component, molecular function, and biological process. Percentage and number of genes assigned to each GO term are indicated.

The sequencing data, and data mapping statistics are provided in Supplementary Tables 3-5. All samples showed high sequencing depth, ranging from 41.7 to 53.1 million total reads per library. Alignment rates were consistently high. Unique mapped reads accounted for >95 % of total reads across samples, further confirming mapping quality. Totally, 25,834 gene expression profile was identified (Supplementary File 1). The principal component analysis (PCA) and correlation heatmap were performed on FPKM of each sample (Fig. 3B-C). Three different biological samples (n=1-3) from each experimental group share similar transcriptomic profiling. The PCA plot shows that samples from the control group (C1–C3) cluster together and are clearly separated from the doxorubicin-treated group (Dx1–Dx3) along PC1, which explains 62.74% of the total variance, while PC2 accounts for 13.81%. This distinct clustering indicates that Dx treatment induces substantial transcriptional changes in human microglial HMC3 cells. The sample-to-sample correlation heatmap further confirms the consistency of the RNA-seq data. High correlation coefficients were observed among biological replicates within the same group, with control samples showing correlations of approximately 0.992 – 0.994 and Dx-treated samples showing correlations of approximately 0.988–0.990. In contrast, lower correlation values (∼0.880–0.889) were observed between control and Dx-treated samples. The hierarchical clustering analysis also grouped samples according to their treatment conditions, separating control and Dx-treated samples into two distinct clusters. Together, these results indicate high reproducibility among biological replicates and a clear transcriptional distinction between control and Dx-treated microglial cells, confirming that the RNA-seq data are reliable for downstream differential gene expression and pathway analyses.

Gene Ontology (GO) classification was performed to functionally annotate all identified genes (Fig.3D and Supplementary File 2). Within the cellular component category, the majority of genes were associated with cellular anatomical entities and intracellular components, with substantial representation in protein-containing complexes. In the molecular function category, genes were predominantly enriched in binding and catalytic activities. Notably, antioxidant activity and molecular function regulator activity were also identified, suggesting the involvement of genes in redox homeostasis and regulatory control mechanisms. Regarding the biological process domain, the identified genes were mainly involved in cellular processes, metabolic processes, and biological regulation. Importantly, immune system process and response to stimulus were evident, indicating activation of stress-responsive and immune-related pathways.

Furthermore, gene set enrichment analysis (GSEA) based on KEGG pathway annotation identified the two most significantly enriched pathways following doxorubicin treatment (Fig. 4A-B and Supplementary File 3). Specifically, the cytokine–cytokine receptor interaction pathway (NES = 1.634, p = 0.001017) and the neuroactive ligand–receptor interaction pathway (NES = 1.661, p = 0.001016) showed strong enrichment, highlighting the coordinated upregulation of immune-related and neuroactive pathways under the doxorubicin-treated condition.

**Figure 4.**
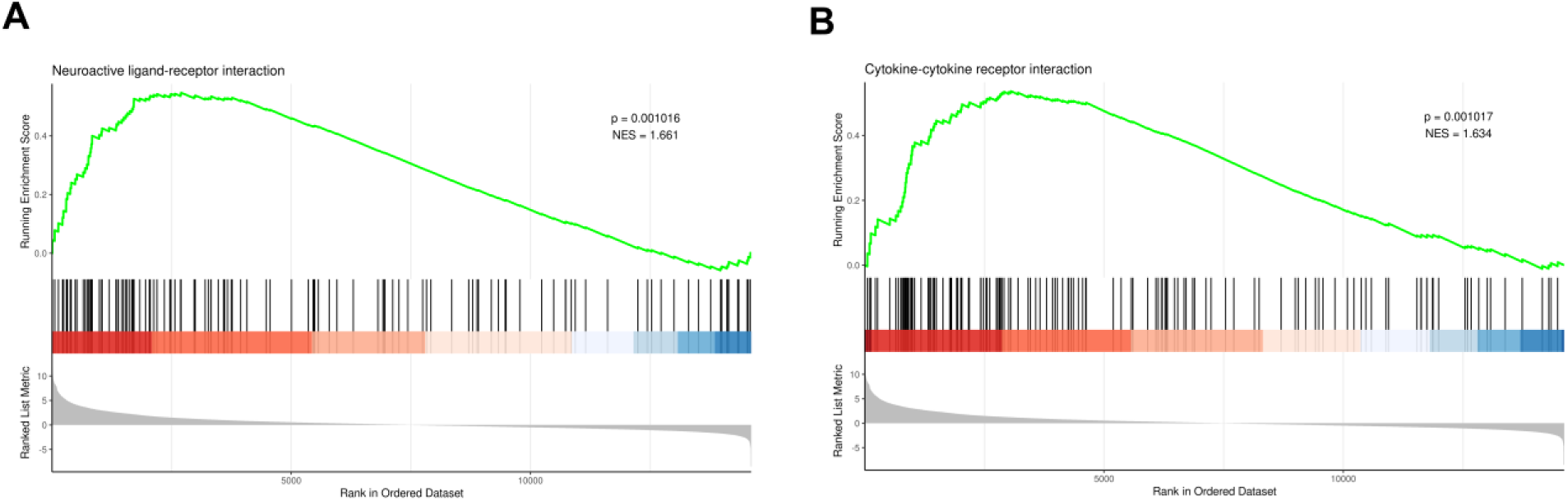
Gene set enrichment analysis (GSEA) reveals the distribution of genes associated with the cytokine–cytokine receptor interaction and neuroactive ligand–receptor interaction pathways. GSEA was performed using KEGG pathway annotation to identify significantly enriched pathways in the transcriptomic dataset (control vs doxorubicin). Enrichment plots for the cytokine–cytokine receptor interaction (**A**) and neuroactive ligand–receptor interaction (**B**) pathways are shown. The green curve represents the running enrichment score (ES), black vertical bars indicate the positions of pathway-associated genes within the ranked gene list, and the color gradient reflects the correlation of gene expression with the control (blue) or doxorubicin (red) condition. Both pathways showed significant enrichment (p<0.05). Full list provided in Supplementary File 3.

### 3.4 A large number of genes were differentially expressed in HMC3 microglial cells following doxorubicin treatment

The list of differentially expressed genes (DEGs) in doxorubicin-treated cells compared to control was analyzed by DESeq2. Criteria for DEGs was set as fold change (FC)≥2 and FDR<0.01. The DEGs were identified and are summarized in Supplementary File 4. In total, 5,669 DEGs were detected between doxorubicin and control. In doxorubicin - treated cells, a large number of genes were upregulated in response to cellular stress, as reflected by the higher number of upregulated DEGs (3,431) compared to downregulated DEGs (2,238) (Fig. 5A). The higher proportion of upregulated DEGs following doxorubicin treatment indicates that the drug primarily activates stress-response and damage-associated transcriptional pathways in human microglial cells. MA plots and volcano plots visualize the overall distribution of gene expression and fold change of expression level between two comparing samples (Fig. 5 B-C). Together with heatmap, it revealed a clear distinction between comparative groups based on their abundance profile of DEGs (Fig. 5D).

**Figure 5.**
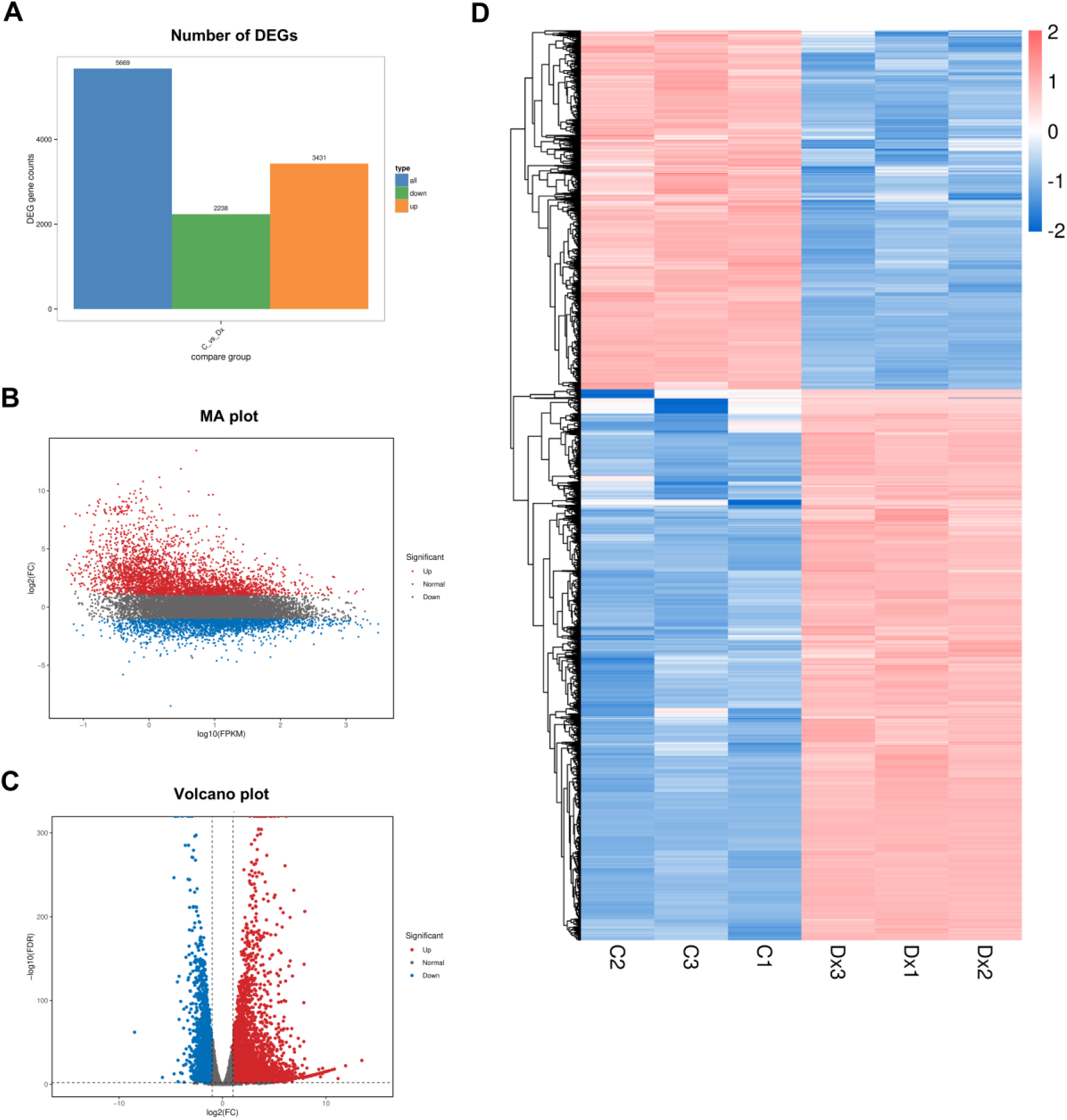
A large number of differentially expressed genes (DEGs) under doxorubicin treatment in human microglial HMC3 cells. **(A)** Total number of DEGs between doxorubicin (Dx) vs control (C). Blue bar represents a total number of DEGs. Green and orange bars indicate the number of DEGs that were downregulated and upregulated under Dx treatment. Criteria for DEGs was set as fold change (FC)≥2 and FDR<0.01. **(B)** MA plot of DEGs illustrates the overall distribution of gene expression and fold change of expression level between two groups. Each dot represents a single gene. X-axis represents values of log2 (FPKM). Y-axis represents values of log_2_(FC). **(C)** Volcano plot visualizes the DEGs in the comparing groups. X-axis represents log2(Fold change) of expression. Y-axis represents -log_10_(FDR). The dots colored in red and blue stand for significant up-regulated and down-regulated genes respectively. Gray dots are genes without significant difference. **(D)** Heatmaps representing hierarchical clustering of DEGs between doxorubicin and control group, based on the expression profile of DEGs. Red color and blue color indicate up-and down-regulated levels of DEGs, respectively.

KEGG pathway enrichment analysis of all DEGs revealed that the differentially expressed genes were significantly involved in pathways related to the ribosome, p53 signaling pathway, apoptosis, base excision repair, and regulation of the actin cytoskeleton (Fig. 6A). These pathways are consistent with our observations that doxorubicin treatment induces DNA damage and cellular senescence, and promotes morphological alterations in the human microglial HMC3 cells. Circular network diagram illustrating gene-function associations of total DEGs (Fig. 6B). Genes are arranged around the circle and connected to their respective KEGG terms. This visualization highlights p53 signaling pathways and ribosome playing a key role cellular response to doxorubicin-induced genotoxic stress response in the HMC3 cells. Additionally, based on GO enrichment analysis, the biological processes associated with doxorubicin-treated cells can be broadly categorized into three groups. The first group is related to translation (highlighted in red). The second group includes processes associated with microtubule-based movement, epithelial morphogenesis, microtubule-based processes, microtubule cytoskeleton organization, cellular localization, positive regulation of epithelial-to-mesenchymal transition, and epithelial tube morphogenesis (highlighted in blue). The third group involves the regulation of cyclin-dependent protein kinase activity (highlighted in green), which may underlie the cell-cycle arrest and senescence observed in the doxorubicin-treated microglial cells (Fig. 6C).

**Figure 6.**
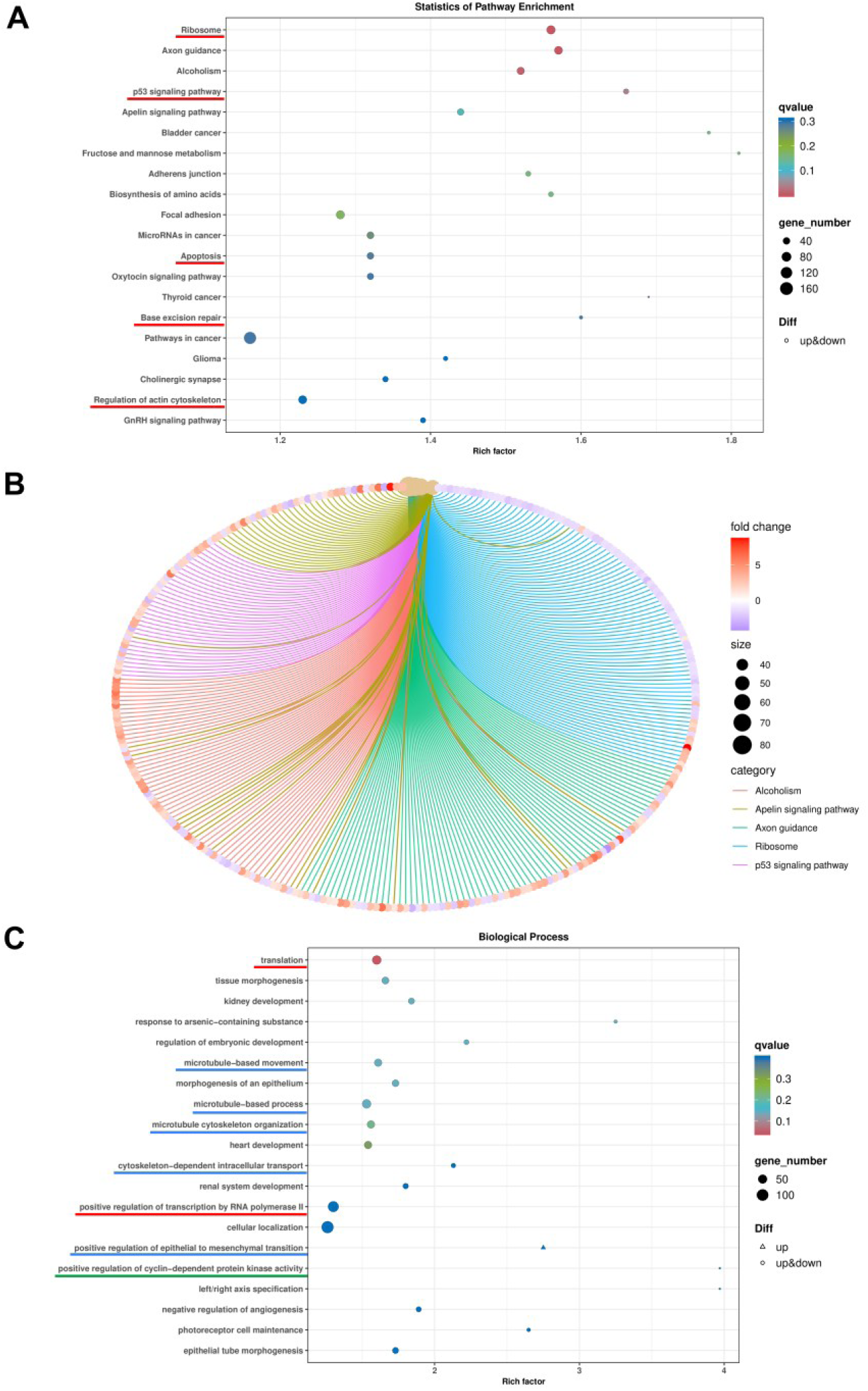
The differentially expressed genes were significantly involved in pathways related to the p53 signaling pathway, apoptosis, base excision repair, gene expression regulation, and cytoskeleton-associated processes. **(A)** KEGG pathway enrichment analysis of total DEGs, showing the enriched pathways associated with doxorubicin-induced genotoxic stress in the HMC3 cells. **(B)** Circular network diagram illustrating gene-function associations of total DEGs. Genes are arranged around the circle and connected to their respective KEGG terms. Node color represents fold change (from blue to red), and black circles indicate gene counts per term. This visualization highlights DEGs playing a key role to p53 signaling pathways underlying the effect of doxorubicin toxicity in human microglial cells. **(C)** GO enrichment analysis of total DEGs. The scatter plot displays enriched biological processes (BP) based on the Rich factor (x-axis), with the number of genes involved represented by the size of each dot and the q-value indicated by color (from blue to red, where red denotes higher significance).

### 3.5 Genes associated to p53 signaling and inflammation mediators were upregulated in doxorubicin-treated HMC3 cells

Next, to identify pathways associated with the upregulated genes following doxorubicin treatment, enrichment analyses using KEGG and GO were performed. The output is provided in Supplementary File 5. Importantly, KEGG pathway analysis revealed that doxorubicin treatment significantly upregulated genes associated with the p53 signaling pathway, regulation of TRP channels by inflammatory mediators, and the TGF-β signaling pathway (Fig. 7A). These pathways are closely associated with DNA damage response, inflammatory signaling, and cellular stress adaptation, suggesting that doxorubicin exposure activates multiple molecular mechanisms related to genotoxic stress, and inflammation in microglial HMC3 cells.

**Figure 7.**
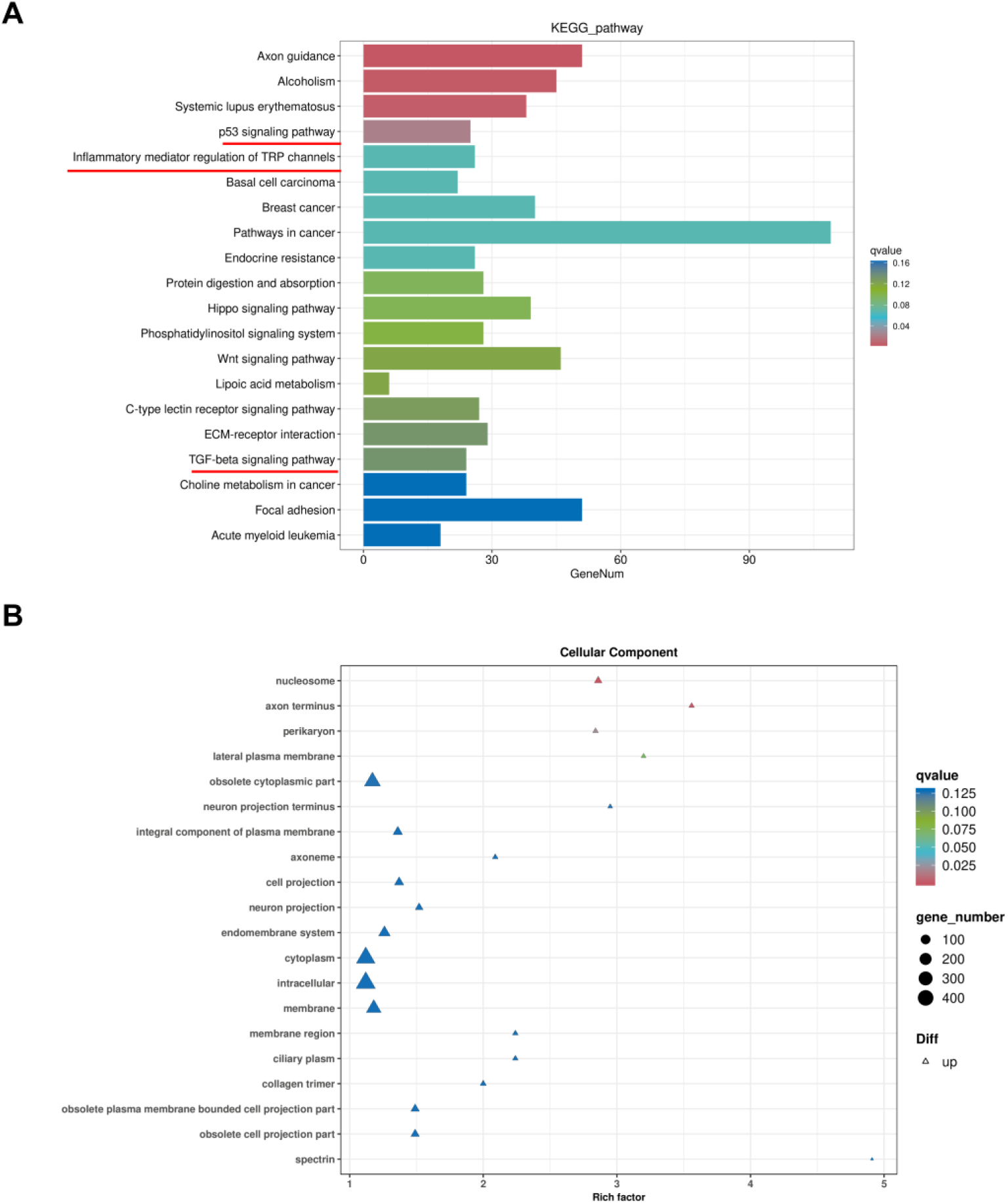
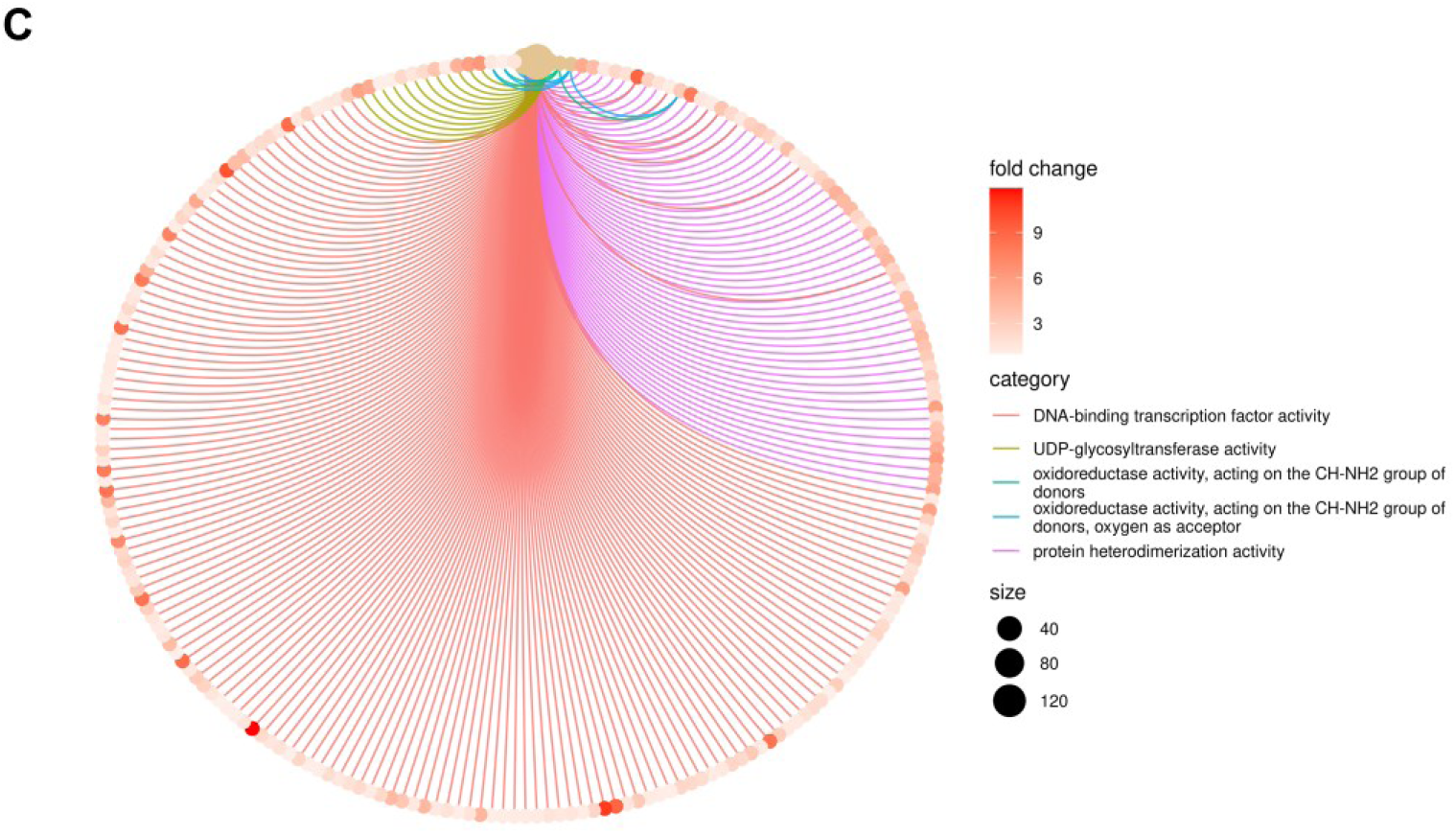
Doxorubicin treatment upregulated genes involved in the p53 signaling pathway, inflammatory mediator regulation of TRP channels, and the TGF-β signaling pathway. **(A)** KEGG pathway enrichment analysis of upregulated DEGs in doxorubicin vs control. Bar chart showing the number of DEGs associated with significantly upregulated KEGG pathways. Bars are color-coded by q-value, and the x-axis represents the number of genes involved in each pathway. **(B)** GO enrichment analysis of upregulated DEGs. The scatter plot displays enriched cellular compartments (CC) based on the Rich factor (x-axis), with the number of genes involved represented by the size of each dot and the q-value indicated by color (from blue to red, where red denotes higher significance). **(C)** Circular network diagram illustrating gene-function associations of molecular function (MF) of the upregulated DEGs. This visualization highlights upregulated DEGs associated with DNA-binding transcription factor activity in response to doxorubicin exposure in human microglial cells.

GO enrichment analysis of the cellular component indicates that a large proportion of upregulated genes are associated with intracellular, cytoplasmic, and membrane-related components, reflecting widespread cellular responses to doxorubicin treatment (Fig. 7B). Notably, the enrichment of the nucleosome category is particularly important, as it suggests alterations in chromatin organization and histone-associated processes, which are closely linked to DNA damage responses and repair mechanisms.

The circular network visualizing molecular function associated to the upregulated genes illustrates that, among the enriched terms, DNA-binding transcription factor activity represents the most prominent category, indicating robust activation of transcriptional regulators (Fig. 7C). This suggests extensive transcriptional reprogramming as a central response to doxorubicin-induced stress. In addition, oxidoreductase activities, particularly those acting on CH–NH2 groups, were also significantly represented, highlighting redox-related processes and oxidative stress responses.

The list of genes and their expression profile compared in two groups (doxorubicin vs control) are presented in Figure 8A-D. Gene annotated to P53 signaling pathway, Inflammatory mediator regulation of TRP channels, TGF-beta signaling pathway, and cytokine-cytokine receptor interaction are listed. Collectively, these genes were upregulated following doxorubicin treatment, reflecting transcriptional alterations associated with genotoxic stress responses.

**Figure 8.**
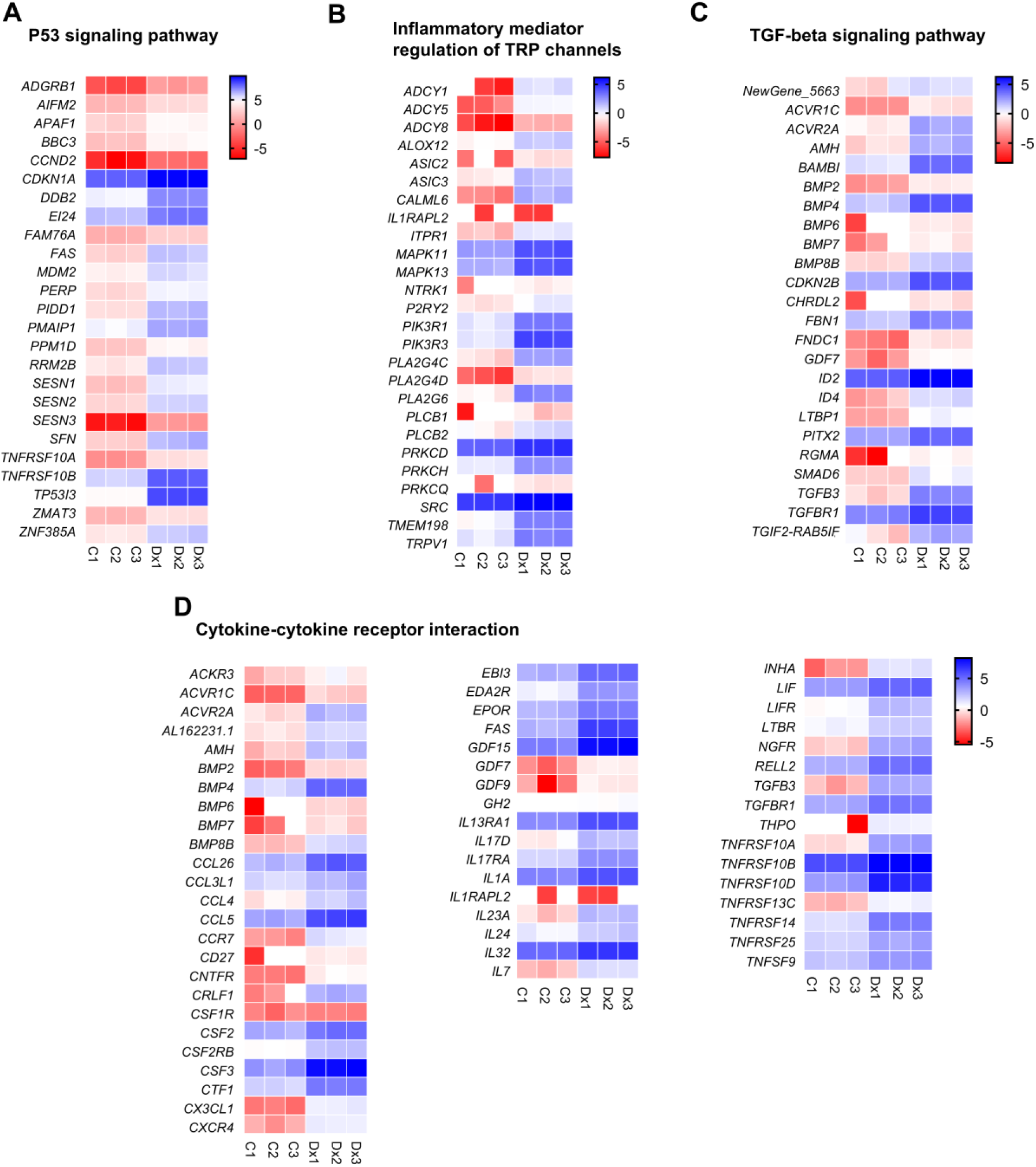
Doxorubicin treatment induced the upregulation of genes associated with the (A) p53 signaling pathway, (B) inflammatory mediator regulation of TRP channels, (C) TGF-β signaling pathway, and (D) cytokine–cytokine receptor interaction in HMC3 microglial cells. Gene expression levels are presented as log_2_ (FPKM) values from three independent biological replicates (n1-3).

### 3.6 Transcriptional suppression of ribosomal and translational genes is consistent with the experimental observation of a reduced total protein and cell cycle shutdown in doxorubicin-treated microglia cells

The downregulated DEGs were annotated using COG functional classification of Consensus sequence (Fig. 9A). The finding suggests that doxorubicin treatment leads to a suppression of the cellular protein synthesis machinery. The enrichment of downregulated DEGs in the translation, ribosomal structure, and biogenesis (COG category J) indicates that genes responsible for ribosome function and protein production are significantly reduced in expression. As shown in Figure 9B, the circular plot demonstrates that doxorubicin-induced transcriptional alteration is not restricted to a single cellular compartment but downregulate genes distributed across multiple intracellular organelles. The largest proportion of downregulated genes is associated with the nucleoplasm, indicating that nuclear processes involved in transcriptional regulation, DNA replication, chromatin remodeling, and RNA processing are broadly inhibited following doxorubicin treatment. A substantial number of downregulated genes are also localized to the mitochondrion, suggesting impaired mitochondrial function. Another prominent group of suppressed genes is associated with the ribosome, indicating reduced protein synthesis. Downregulation of ribosomal components is commonly observed during cellular stress, where global translation is attenuated to conserve energy while selectively translating stress-response proteins. Although fewer in number, spindle-associated genes are also downregulated, implying inhibition of mitotic progression and cell division, thereby preventing mitotic entry.

**Figure 9.**
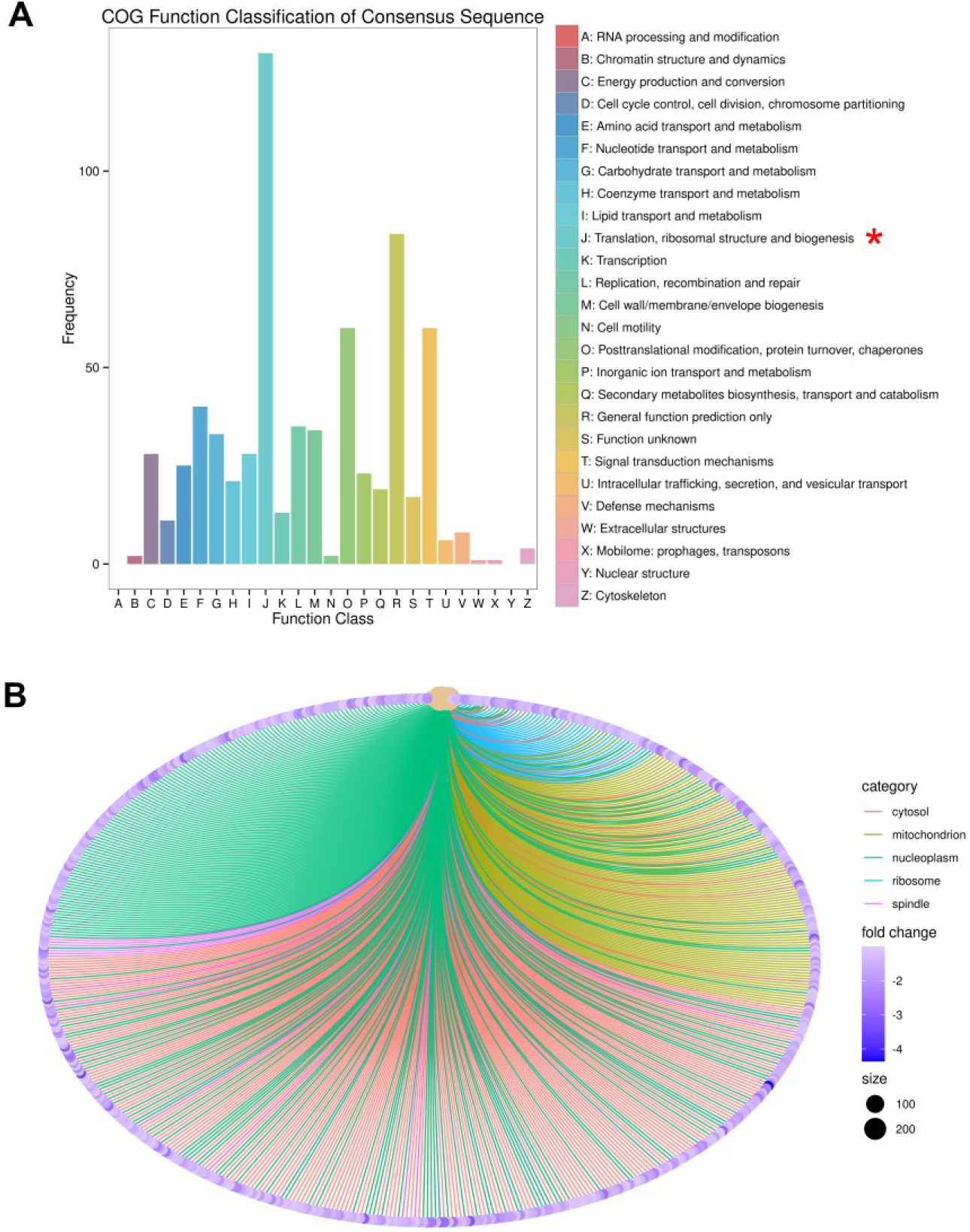
Doxorubicin treatment suppressed the cellular protein synthesis machinery, as indicated by the downregulation of genes involved in translation and ribosome-related functions. **(A)** The eggNOG functional classification of consensus sequences of downregulated DEGs. The bar chart displays the frequency of sequences assigned to various functional categories (A–Z) based on eggNOG annotations. Notably, the downregulated DEGs in doxorubicin are enriched in translation, ribosomal structure and biogenies (category J). **(B)** Circular network diagram illustrating gene-function associations of cellular compartment (CC) of the downregulated DEGs. The visualization highlights the downregulation of genes associated with ribosomal and nucleosomal components in response to doxorubicin exposure in human microglial cells.

Furthermore, KEGG pathway analysis revealed that genes involved in the ribosome, biosynthesis of amino acids, nucleotide excision repair, DNA replication, homologous recombination, cell cycle, and base excision repair were significantly downregulated in doxorubicin-treated microglial HMC3 cells (Fig.10A). GO enrichment analysis further confirmed that genes associated with translation and cell division were suppressed in doxorubicin-treated cells (Fig.10B). These molecular changes in gene expression are consistent with the phenotypic observations, indicating impaired cell-cycle progression in response to doxorubicin-induced genotoxic stress. Transcriptional suppression of ribosomal and translational genes is also consistent with the experimental observation of a reduced total protein yield in doxorubicin-treated cells, suggesting that genotoxic stress induced by doxorubicin impairs the protein synthesis capacity of microglia (Supplementary Fig. 1). Overall, doxorubicin induces translational repression commonly associated with cellular stress responses, DNA damage signaling, and the transition toward cell-cycle arrest or senescence, where cells reduce energy-demanding processes such as protein production to maintain cellular homeostasis under stress conditions.

### 3.7 Validation of differentially expressed genes underlying doxorubicin-induced genotoxic stress in HMC3 cells

The mRNA-seq analysis identified several genes that were upregulated under Dx-induced genotoxic stress in human microglia cells. Therefore, it is necessary to validate the expression of these genes to confirm the reliability of the dataset. We validated genes including anti-oxidant genes (*NFE2L2*, *GPX*, *SOD1*, and *CAT)*, senescence-or apoptosis-related genes (*TP53*, and *P21*), and stress-responsive gene (*GDF15, SIRT1*) using RT-PCR.

The gene expression levels from real-time PCR (Fig.10 A-H) and the correlation of the expression profiles obtained from the two methods (Fig.10I) is presented. The analysis demonstrates a significant positive correlation between the two methods (r = 0.79, R ^2^= 0.6247, p = 0.0196), confirming the reliability and consistency of the RNA-seq data with RT-PCR validation.

**Figure 10.**
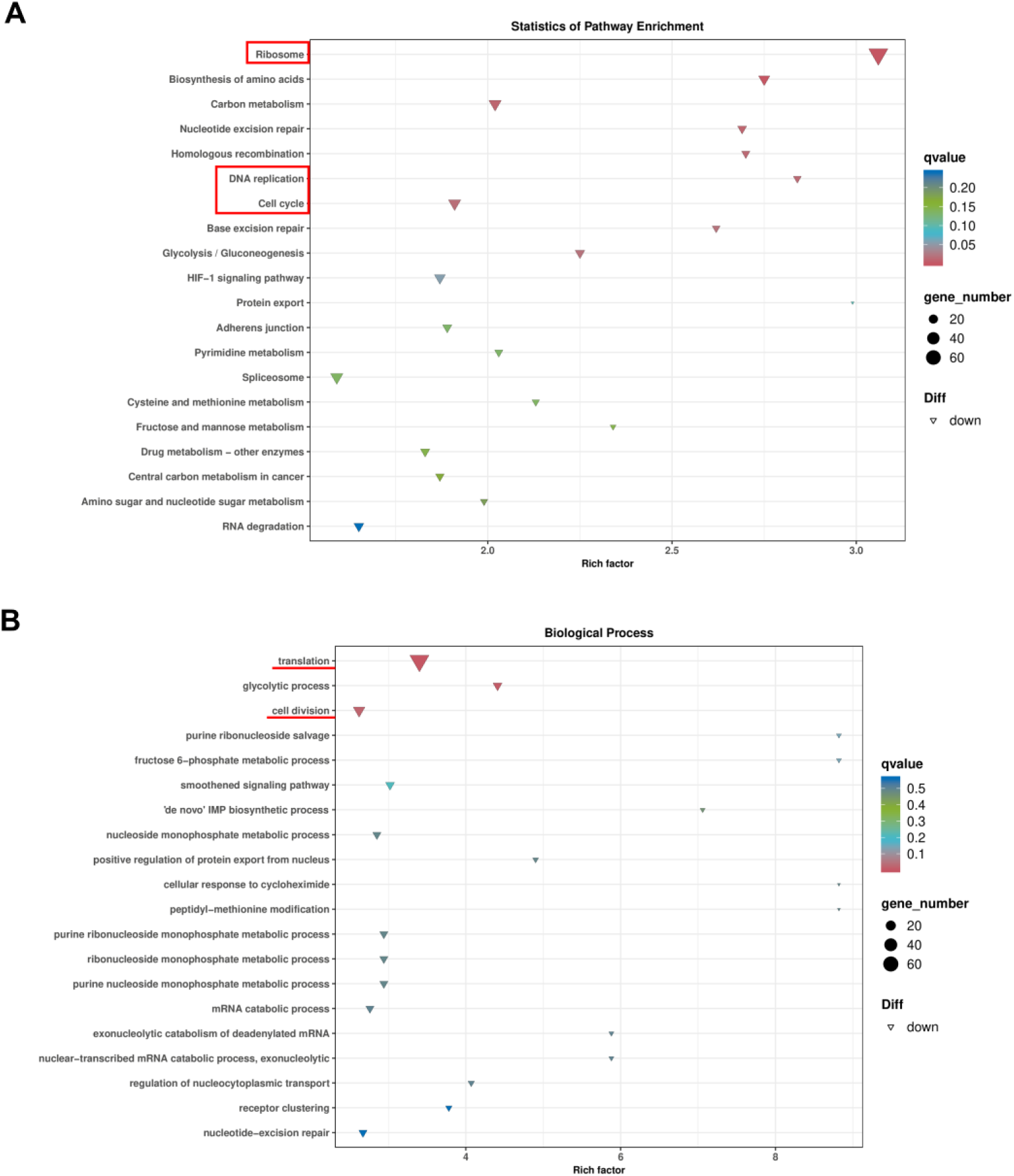
KEGG and GO enrichment analyses revealed significant downregulation of genes involved in ribosome biogenesis and translation, DNA replication, cell cycle regulation, and base excision repair in doxorubicin-treated human microglial HMC3 cells. **(A)** KEGG pathway enrichment analysis of downregulated DEGs in doxorubicin. Bar chart showing the number of DEGs associated with significantly downregulated KEGG pathways. Bars are color-coded by q-value, and the x-axis represents the number of genes involved in each pathway. **(B)** GO enrichment analysis of downregulated DEGs. Bar chart showing the number of genes associated with significantly downregulated biological processes (BP). Bars are color-coded by q-value, and the x-axis represents the number of genes involved in each BP. Genes involving translation, and cell division were highlighted.

### 3.8 Transcriptional factors

Transcriptional factors (TFs) play a vital role in regulating cellular stress responses. To extensively explore TFs involved in doxorubicin -regulated microglial transcriptomic changes, TFs of DEGs were identified and annotation based on AminalTFDB database (Supplementary File 6). As shown in Figure 11, doxorubicin substantially altered the TFs landscape in human HMC3 microglial cells. A total of 414 differentially TFs were identified and classified by transcriptional factor family (Fig.11A). The most prominent family was C2H2 zinc-finger (179 TFs), followed by 36 TFs from Homeobox, 28 TFs from bHLH, and 19 TFs from TF_bZIP.

**Figure 11.**
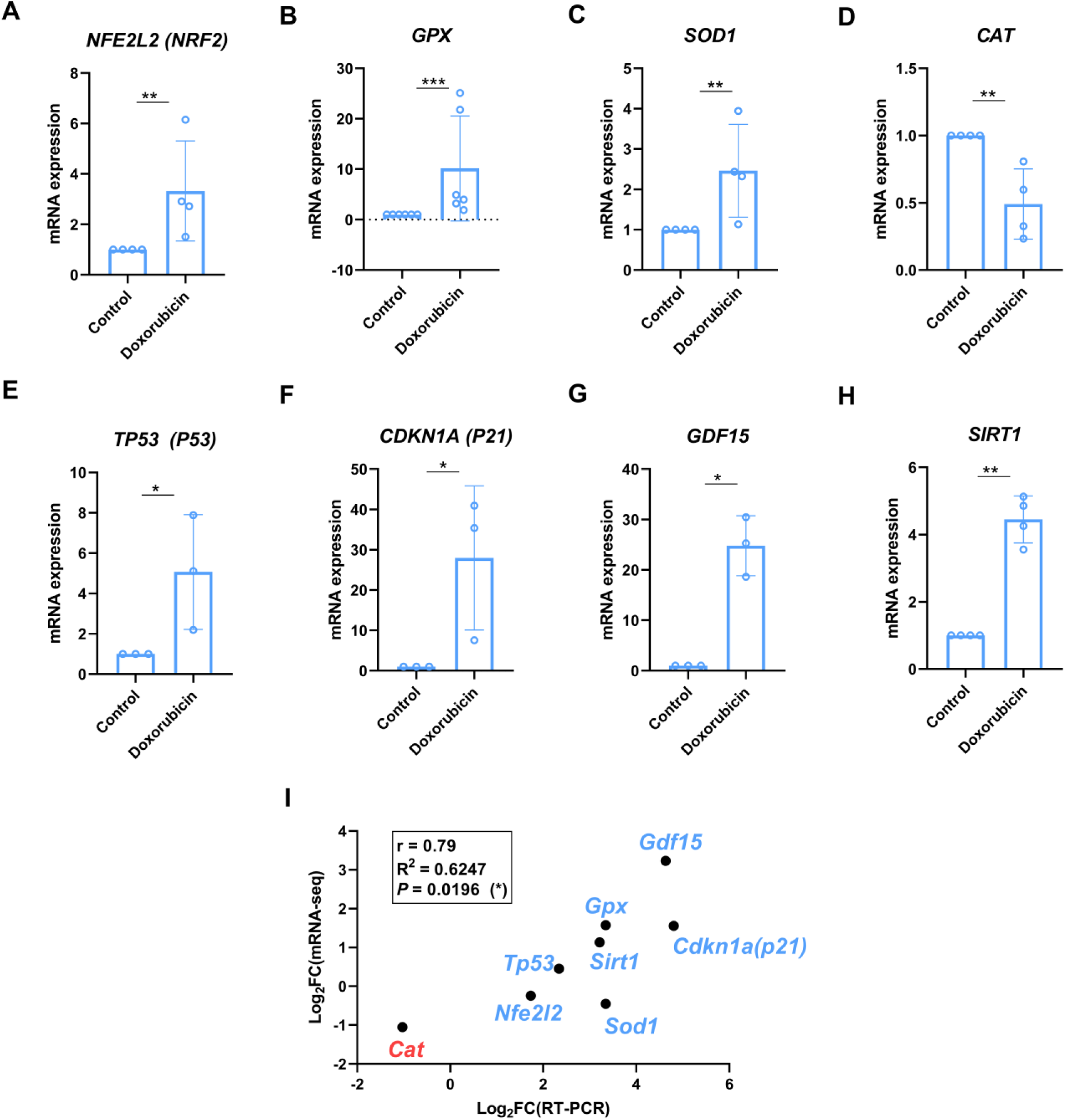
Positive correlation between gene expression profiles obtained by RT-PCR and mRNA-seq. Genes associated with antioxidant activity **(A–D)**, p53 signaling **(E–F)**, and stress responses **(G–H)** were analyzed by RT-PCR. **(I)** Pearson correlation analysis demonstrating the concordance between gene expression profiles obtained by RT-PCR and mRNA-seq.

**Figure 12.**
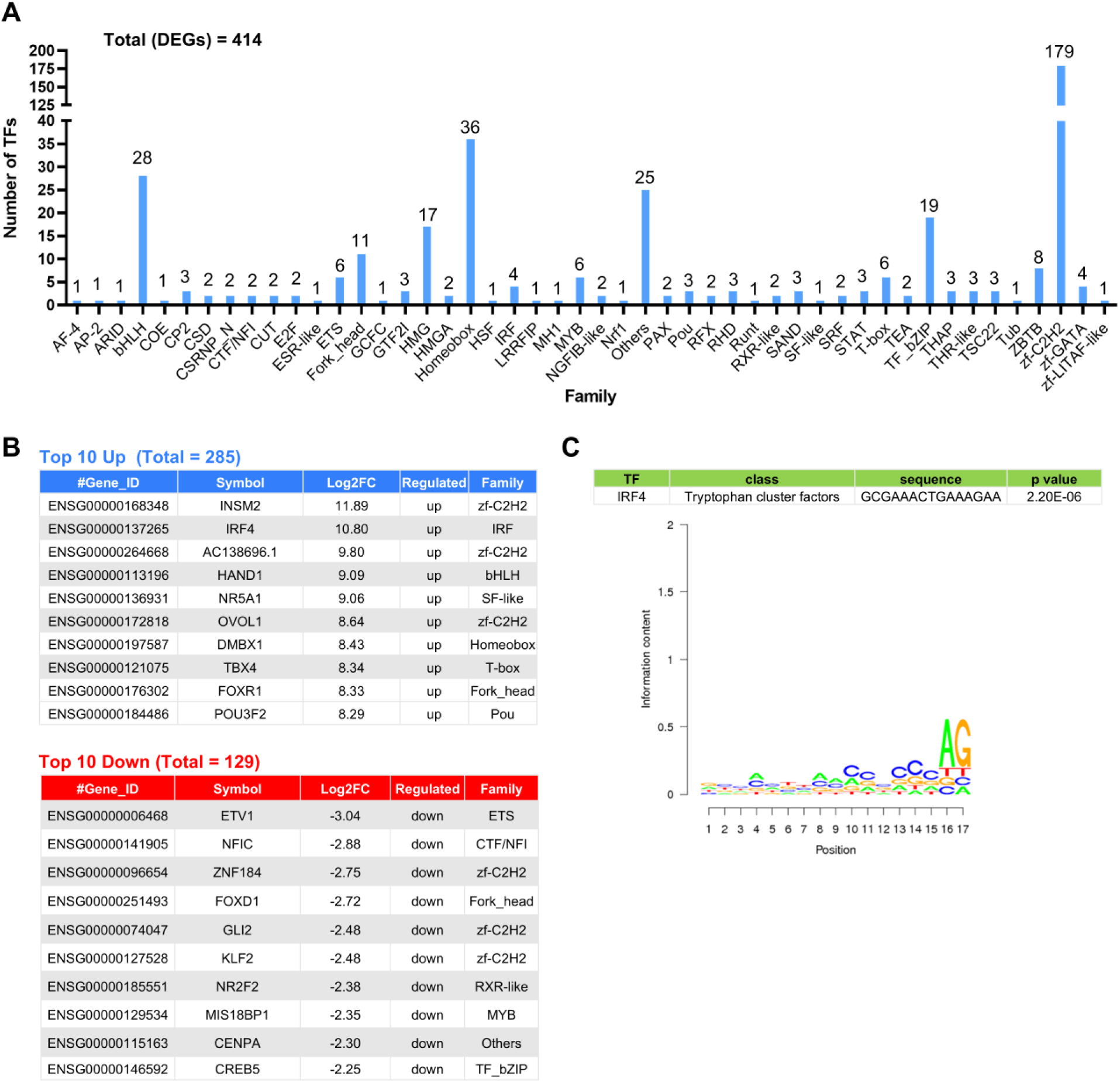
Transcriptional factors (TF) were differentially expressed under doxorubicin treatment in human microglial HMC3 cells. **(A)** Category of 414 TFs that were differentially expressed under doxorubicin treatment in human microglial HMC3 cells. **(B)** Top TFs altered by doxorubicin treatment. The data present the Gene ID, gene symbol, log_2_FC, regulation status (up or down), and TF family classification. **(C)** The predicted transcription factor binding site (TFBS) analysis for IRF4 (Interferon Regulatory Factor 4). The predicted TFBS information was obtained using the TFBSTools package with the JASPAR motif database. The sequence logo plot represents the consensus binding motif of IRF4, where the height of each nucleotide reflects its information content and conservation across the motif positions. Highly conserved bases indicate key nucleotides required for IRF4 binding specificity.

Of these, 285 TFs were upregulated, whereas 129 were downregulated, indicating a predominantly inductive transcriptional response to doxorubicin (Fig. 11B). Among the upregulated TFs, INSM2 showed the largest increase, followed by IRF4, AC138696.1, HAND1, NR5A1, OVOL1, DMBX1, TBX4, FOXR1, and POU3F2. Conversely, the most strongly downregulated TFs included ETV1, NFIC, ZNF184, FOXD, GLI2, KLF2, NR2F2, MIS18BP1, CENPA, and CREB5.

One particularly notable finding is IRF4, which was among the most strongly induced TFs. IRF4 is a transcriptional regulator involved in cellular differentiation and immune-related transcriptional programs. A significant IRF4-associated DNA-binding motif was identified (Fig. 11C), providing computational evidence that IRF4 -associated transcriptional regulation may contribute to the doxorubicin-induced transcriptional response. Further studies are needed to determine whether targeting the transcriptional activity of IRF4 can mitigate the deleterious effects induced by doxorubicin.

## 4. Discussion

Chemotherapy-related cognitive impairment is increasingly recognized as an important complication of cancer treatment, yet the cellular mechanisms underlying chemotherapy-associated neurotoxicity remain unresolved. Although previous studies have largely focused on neuronal responses to chemotherapeutic agents, accumulating evidence indicates that non-neuronal cells of the CNS, particularly microglia, also play important roles in chemotherapy-associated neurotoxicity. In the present study, the cellular and transcriptomic responses of human HMC3 microglial cells to doxorubicin-induced genotoxic stress were investigated. Our findings demonstrate that doxorubicin induces persistent growth inhibition, morphological alterations, DNA damage, and senescence-associated characteristics in HMC3 cells. These phenotypic changes were accompanied by extensive transcriptional reprogramming involving DNA damage responses, p53 signaling, inflammatory pathways, translation, DNA replication, and cell-cycle regulation.

Exposure to doxorubicin (1-5 μM) reduced HMC3 cell viability by approximately 20%, while removal of doxorubicin followed by culture in fresh medium resulted in a more pronounced reduction in viability. This observation suggests that doxorubicin induced sustained cellular stress rather than a transient reduction in metabolic activity. Doxorubicin treatment induced pronounced morphological changes in HMC3 cells, including an elongated, process-bearing morphology and increased Iba1 intensity. These morphological and immunocytochemical changes are indicative of an altered microglial state and are consistent with previous reports describing microglial activation to doxorubicin exposure(Demby et al., 2024; Jia et al., 2023; Moretti et al., 2021).

γH2AX is widely used as a sensitive marker of DNA double-strand breaks and is therefore commonly employed as an indicator of DNA damage(Hosking et al., 2024; Prabhu et al., 2024). Doxorubicin induces DNA damage across diverse cell types, including astrocytes, the most abundant glial cell type in the CNS(Marques et al., 2020). Our recent study demonstrated a significant increase in nuclear γH2AX following doxorubicin exposure in human microglial cells, providing evidence of DNA damage in response to doxorubicin-induced genotoxic stress. Sirtuin-1 (SIRT1) is an NAD⁺-dependent deacetylase that participates in the regulation of genome stability, DNA repair, and stress responses(Rasti et al., 2023). The concomitant increase in SIRT1 may represent an adaptive response to cellular stress following doxorubicin treatment, potentially contributing to the regulation of H2AX phosphorylation, as previously reported in cardiomyocytes(Kuno et al., 2022). Nevertheless, prolonged exposure to higher doses of doxorubicin is associated with reduced SIRT1 expression, suggesting that disruption of SIRT1-mediated stress responses may contribute to increased cellular damage (Ebrahim et al., 2024).

The reduction in Ki67-positive cells and increase in senescence-associated β-galactosidase-positive cells further support the development of a senescence-associated phenotype after doxorubicin exposure in the HCM3 cells. These findings are consistent with previous reports showing that doxorubicin can induce senescence in microglia and other CNS cell populations(Demby et al., 2024; Marques et al., 2020). In particular, doxorubicin-induced senescent microglia have previously been reported to exhibit altered protein profiles, increased cytokine secretion, and impaired amyloid-β internalization, suggesting that senescence may have functional consequences extending beyond proliferative arrest(Marques et al., 2020). Thus, the present findings extend previous observations by demonstrating that human HMC3 microglia exhibit coordinated DNA damage and senescence-associated responses following doxorubicin exposure.

The transcriptomic analysis provides a genome-wide view of the molecular response underlying the observed cellular phenotype. A total of 25,834 expressed genes were detected, of which 5,669 were differentially expressed between control and doxorubicin-treated microglia cells. Notably, 3,431 genes were upregulated compared with 2,238 downregulated genes, suggesting that the response is characterized predominantly by induction of stress- and damage-associated transcriptional programs. Rather than affecting a single molecular pathway, these mechanisms are expected to activate interconnected programs controlling DNA damage sensing, cell-cycle progression, apoptosis, metabolism, inflammatory signaling, and cellular adaptation. These findings are particularly relevant to the potential involvement of microglia in chemotherapy-associated neurological dysfunction. Microglia communicate extensively with neurons and other CNS cells through cytokines, chemokines, growth factors, and other signaling molecules(Di Pietro et al., 2026; Zong et al., 2026). Therefore, transcriptional alterations affecting cytokine-receptor networks could modify microglial communication with neighboring neural cells. However, because the present study measured transcript abundance rather than proteins or functional signaling, these pathway enrichments should be regarded as molecular signatures requiring further functional validation.

One of the most prominent findings was the enrichment of the p53 signaling pathway among genes differentially expressed following doxorubicin treatment. This observation is consistent with the experimental evidence of DNA damage and cell-cycle arrest. Activation of the DNA damage response can engage p53-dependent transcriptional programs that coordinate cell-cycle arrest, DNA repair, senescence, and, depending on the severity of damage, apoptosis. Microglial apoptosis is prevented by pifithrin-α, an inhibitor of the tumor suppressor protein p53 (Davenport et al., 2010). The present data therefore points that direct targeting p53-associated signaling represents an important strategic approach to ameliorate the molecular response to doxorubicin-induced genotoxic stress.

In contrast to the activation of stress-responsive pathways, genes involved in translation, ribosomal function, DNA replication, nucleotide metabolism, homologous recombination, and cell-cycle progression were broadly suppressed. This transcriptional pattern provides a molecular explanation for the observed reduction in cellular proliferation and senescence-associated phenotype. Downregulation of ribosomal and translational machinery may represent a mechanism by which stressed cells reduce energy - demanding biosynthetic processes while prioritizing stress adaptation and survival. The reduction in total protein yield observed experimentally is also consistent with this transcriptional suppression (Supplementary Fig. 1). The decline in total protein is consistent with a previous SWATH-MS-based proteomic study of doxorubicin-exposed immortalized rat microglial CHME-5 cells, which identified a greater number of downregulated than upregulated proteins (Marques et al., 2020). Thus, the simultaneous induction of stress-response genes and repression of fundamental biosynthetic and proliferative programs therefore suggests a coordinated transition from a proliferative cellular state toward a stress-adapted, growth-arrested state.

Furthermore, the present transcriptomic data showed enrichment of antioxidant activity-related molecular functions among upregulated genes (Fig. 3), suggesting that microglia activate compensatory mechanisms in response to altered redox conditions. Consistent with this interpretation, the expression of antioxidant-and stress-related genes, including SIRT1, NFE2L2, GPX, SOD1, CAT, and GDF15, was evaluated by RT-qPCR. The significant positive correlation between RNA-seq and RT-qPCR measurements supports the reliability of the transcriptomic dataset. Nevertheless, prolonged or higher-dose doxorubicin exposure may shift microglia from an adaptive stress response toward cellular dysfunction, warranting further investigation under more severe exposure conditions.

A major objective of the transcriptomic analysis was to move beyond the identification of individual DEGs and investigate transcriptional regulators that may coordinate the observed gene expression changes. The analysis identified 414 differentially expressed transcription factors. Among these, IRF4 was one of the most strongly induced transcription factors, suggesting that IRF4-associated transcriptional activity may represent a potential regulatory component of the microglial response to doxorubicin. IRF4 is a transcriptional regulator with established roles in immune-cell differentiation and regulation of immune-associated gene programs (Ngwa et al., 2024). Direct links connecting IRF4 to microglial DNA damage responses remain largely unexplored in current literature. Its strong induction in doxorubicin-treated HMC3 cells raises the possibility that IRF4 contributes to the transcriptional remodeling of microglia under genotoxic stress. The identification of IRF4 nevertheless provides a potentially valuable direction for subsequent mechanistic studies. Loss-of-function approaches, such as IRF4 knockdown or inhibition, followed by assessment of DNA damage, senescence, inflammatory mediator production, and transcriptomic responses could determine whether IRF4 is required for specific components of the doxorubicin response.

The findings of this study provide a potential cellular framework linking chemotherapy-induced genotoxic stress in microglia to alterations in the CNS microenvironment. Chemotherapy-associated cognitive impairment may arise from interactions among multiple CNS cell types rather than from direct neuronal toxicity alone. Previous studies have demonstrated that doxorubicin can induce brain senescence and that microglial responses may be associated with altered cytokine secretion and impaired cellular functions(Demby et al., 2024; Marques et al., 2020). The present study adds a human microglial transcriptomic perspective and identifies specific molecular pathways and candidate transcriptional regulators associated with the response. However, caution is required when extrapolating the current findings to chemobrain. Several limitations should be considered. First, the study used HMC3 cells as an in vitro human microglial model. Validation in primary human microglia or other physiologically relevant microglial models will be necessary to establish the generalizability of the findings. Second, the transcriptomic analysis was performed using bulk RNA-seq with three biological replicates per condition. Although the high intra-group correlation and clear separation between experimental groups support the robustness of the dataset, additional biological replicates and independent validation would strengthen the statistical confidence of individual gene-level observations. Third, RNA-seq primarily provides information regarding transcriptional regulation and does not necessarily predict protein abundance or cellular function. Future studies should therefore integrate transcriptomic data with complementary omics approaches, including proteomics and metabolomics, to determine whether transcriptional changes are translated into alterations in protein expression and cellular metabolism. Such multi-omics integration could provide a more comprehensive molecular representation of doxorubicin-induced microglial dysfunction. Finally, functional validation should include measurement of cytokine and chemokine secretion, ROS production, ATP levels, mitochondrial respiration, and dynamic proliferative capacity. Co-culture experiments with neurons could further determine whether doxorubicin-treated microglia alter neuronal viability, synaptic function, or neuronal inflammatory responses.

## 5. Conclusion

Overall, this study demonstrates that doxorubicin induces extensive and coordinated transcriptional remodeling in human HMC3 microglial cells. These molecular alterations are accompanied by persistent DNA damage and senescence-associated phenotypic changes. Collectively, these findings establish a transcriptomic framework for investigating the contribution of microglial dysfunction to chemotherapy-associated neurotoxicity and provide a foundation for subsequent multi-omics and functional studies.

## Declaration of Generative AI and AI-assisted technologies in the writing process

The author used Curie in order to improve the English during the preparation of this work. After using this tool, the author reviewed and edited the content as needed and take full responsibility for the content of the published article.

## Contributions

EK: Conceptualization, Data curation, Formal analysis, Investigation, Methodology, Validation, Visualization, Writing – original draft, Funding acquisition.

## Acknowledgment

I sincerely thank Dr. Wasutorn Chankamngoen for technical assistance with the agarose gel electrophoresis experiments. I am also grateful to Prof. Dr. Banthit Chetsawang for providing laboratory resources and support. I further acknowledge BMKGENE for providing Illumina-based transcriptomic sequencing services.

## Data availability

Additional data will be made available on request.

