## Supplementary File for "Microglia transcriptomic response to doxorubicin-induced genotoxic stress"

Ekkaphot Khongkla[1*]

[1] Research Center for Neuroscience, Institute of Molecular Biosciences, Mahidol University, Nakhon Pathom 73170, Thailand

**Supplementary Table 1** List of primary antibodies used in this study.

| Antibody | Supplier | CAS number |
|---|---|---|
| Anti-GAPDH, rabbit polyclonal | Sigma-Aldrich | ABS16 |
| Anti-Iba1, rabbit monoclonal | Cell Signaling Technology | 17198 |
| Anti-ki67, rabbit polyclonal | Abcam | Ab15580 |
| Anti-Phospho-γH2A.X-S139, rabbit polyclonal | Cell Signaling Technology | SAB5700329 |
| Anti-SirT1 (D1D7), rabbit monoclonal | Cell Signaling Technology | 9475S |

**Supplementary Table 2** List of human primers used in this study.

| Target | Forward 5' → 3' | Reverse 5' → 3' |
|---|---|---|
| *NFE2L2 (NRF2)* | CACATCCAGTCAGAAACCAGTGG | GGAATGTCTGCGCCAAAAGCTG |
| *GPX* | CAACCAGTTTGGGCATCAG | TTCACCTCGCACTTCTCG |
| *SOD1* | GATGACTTGGGCAAAGGTGG | TACACCACAAGCCAAACGACT |
| *CAT* | CGTGCTGAATGAGGAACAGA | AGTCAGGGTGGACCTCAGTG |
| *TP53 (P53)* | CCGCAGTCAGATCCTAGCG | AATCATCCATTGCTTGGGACG |
| *CDKN1A (P21)* | CCTGTCACTGTCTTGTACCCT | GCGTTTGGAGTGGTAGAAATCT |
| *GDF15* | GGTGAATGGCTCTCAGATG | CACTTCTGGCGTGAGTATC |
| *SIRT1* | TGCGGGAATCCAAAGGATAA | CAGGCAAGATGCTGTTGCA |
| *GAPDH* | GGAGCGAGATCCCTCCAAAAT | GGCTGTTGTCATACTTCTCATGG |

**Supplementary Table 3** Sample information for mRNA-sequencing.

| #ID | read1_name | read2_name |
|---|---|---|
| CTL1 | Unknown_CQ975-001T0001_good_1.fq | Unknown_CQ975-001T0001_good_2.fq |
| CTL2 | Unknown_CQ975-001T0002_good_1.fq | Unknown_CQ975-001T0002_good_2.fq |
| CTL3 | Unknown_CQ975-001T0003_good_1.fq | Unknown_CQ975-001T0003_good_2.fq |
| Dx1 | Unknown_CQ975-001T0004_good_1.fq | Unknown_CQ975-001T0004_good_2.fq |
| Dx2 | Unknown_CQ975-001T0005_good_1.fq | Unknown_CQ975-001T0005_good_2.fq |
| Dx3 | Unknown_CQ975-001T0006_good_1.fq | Unknown_CQ975-001T0006_good_2.fq |

Note: #ID:sample name; read1_name & read2_name:Double-ended fq file name corresponding to the sample name.

**Supplementary Table 4** Sequencing data Statistics.

| #SampleID | ReadSum | BaseSum | GC (%) | N (%) | Q20(%) | Q30(%) |
|---|---|---|---|---|---|---|
| CTL1 | 26566901 | 7900072705 | 49.5 | 0 | 99.35 | 96.68 |
| CTL2 | 24110803 | 7140115023 | 50.43 | 0 | 99.38 | 96.87 |
| CTL3 | 21098951 | 6306158254 | 49.8 | 0 | 99.32 | 96.59 |
| Dx1 | 23102812 | 6908497646 | 49.96 | 0 | 99.3 | 96.39 |
| Dx2 | 20855344 | 6224069719 | 50.29 | 0 | 99.32 | 96.47 |
| Dx3 | 22433612 | 6696118233 | 50.92 | 0 | 99.32 | 96.53 |

Note:
(1) Samples: Sample name;
(2) Clean reads: Counts of clean PE reads;
(3) Clean bases: total base number of Clean Data;
(4) GC content: Percentage of G, C in clean data.
(5) ≥Q30%: Percentage of bases with Q-score no less than Q30.

**Supplementary Table 5** Statistics on data mapping.

| Sample | Total Reads | Mapped Reads | Uniq Mapped Reads | Multiple Map Reads | Reads Map to '+' | Reads Map to '-' |
|---|---|---|---|---|---|---|
| CTL1 | 53,133,802 | 52,159,170 (98.17%) | 50,825,570 (95.66%) | 1,333,600 (2.51%) | 27,058,907 (50.93%) | 27,042,774 (50.90%) |
| CTL2 | 48,221,606 | 47,407,573 (98.31%) | 45,981,424 (95.35%) | 1,426,149 (2.96%) | 24,950,510 (51.74%) | 24,936,861 (51.71%) |
| CTL3 | 42,197,902 | 41,448,642 (98.22%) | 40,388,017 (95.71%) | 1,060,625 (2.51%) | 21,523,700 (51.01%) | 21,505,421 (50.96%) |
| Dx1 | 46,205,624 | 45,178,077 (97.78%) | 44,132,256 (95.51%) | 1,045,821 (2.26%) | 23,393,268 (50.63%) | 23,381,771 (50.60%) |
| Dx2 | 41,710,688 | 40,902,792 (98.06%) | 39,918,941 (95.70%) | 983,851 (2.36%) | 21,215,592 (50.86%) | 21,205,233 (50.84%) |
| Dx3 | 44,867,224 | 44,005,971 (98.08%) | 42,946,353 (95.72%) | 1,059,618 (2.36%) | 22,830,528 (50.88%) | 22,821,208 (50.86%) |

Note:
Sample: sample ID in system;
Total Reads: Counts of Clean Reads, counted as single end;
Mapped Reads: Counts of mapped reads and the proportion of that in clean data;
Uniq Mapped Reads: Counts of reads mapped to a unique position on reference genome and proportion of that in clean data;
Multiple Mapped Reads: Counts of reads mapped to multiple positions on reference genome and proportion of that in clean data;
Reads Map to '+': Counts of reads mapped to the sense chain and the proportion of that in clean data;
Reads Map to '-': Counts of reads mapped to antisense chain and proportion of that in clean data.

**Supplementary Table 6** Software, and database used in this study.

| Name | Description | Linkages |
|---|---|---|
| **Software** | | |
| HISAT2 | A spliced read mapper for RNA-Seq | http://ccb.jhu.edu/software/hisat2/index.shtml |
| StringTie | Transcript assembly for RNA-Seq | https://ccb.jhu.edu/software/stringtie/index.shtml |
| ASprofile | ASprofile is a suite of programs for extracting, quantifying and comparing alternative splicing (AS) events from RNA-seq data | http://ccb.jhu.edu/software/ASprofile/ |
| BLAST | Basic Local Alignment Search Tool | http://blast.ncbi.nlm.nih.gov/Blast.cgi |
| DESeq | An R package for RNA-Seq Differential Expression Analysis based on a model using the negative binomial distribution | http://www.bioconductor.org/packages/release/bioc/html/DESeq.html |
| EBSeq | An R package for RNA-Seq Differential Expression Analysis based on Bayesian approach | https://www.biostat.wisc.edu/~kendzior/EBSEQ/ |
| Cytoscape | An open source software platform for visualizing complex networks | http://www.cytoscape.org/ |
| topGO | An R package for gene ontology enrichment analysis | https://bio.tools/topgo |
| rMATs | MATS is a computational tool to detect differential alternative splicing events from RNA-Seq data. | http://rnaseq-mats.sourceforge.net/ |
| TFBStools | An R package for the analysis and manipulation of transcription factor binding sites. | http://www.bioconductor.org/packages/release/bioc/html/TFBSTools.html |
| **Database** | | |
| NR | non-redundant protein sequence database | ftp://ftp.ncbi.nih.gov/blast/db/ |
| Swiss-Prot | A manually annotated, non-redundant protein sequence database | http://www.uniprot.org/ |
| GO | Gene Ontology database | http://www.geneontology.org/ |
| COG | The database of Clusters of Orthologous Groups of proteins | http://www.ncbi.nlm.nih.gov/COG/ |
| KOG | The database of Clusters of Protein homology | http://www.ncbi.nlm.nih.gov/KOG/ |
| Pfam | The database of Homologous protein family | http://pfam.xfam.org/ |
| KEGG | The database of Kyoto Encyclopedia of Genes and Genomes | http://www.genome.jp/kegg/ |
| STRING | Search Tool for the Retrieval of Interacting Genes/Proteins | http://www.string-db.org/ |
| Ensembl | Database Sscrofa10.2 download from | http://asia.ensembl.org/index.html |
| Cosmic | COSMIC, is the world's largest and most comprehensive resource for exploring the impact of somatic mutations in human cancer. | https://cancer.sanger.ac.uk/cosmic |

| | | |
|---|---|---|
| JASPAR | Database of transcription factor binding profiles | http://jaspar.genereg.net/ |

**Supplementary Figure 1**

Measurement of **(A)** total RNA yield (ng) and **(B)** total protein concentration (µg) isolated from untreated- and doxorubicin (dx, 1µM)-treated cells by Nanodrop and BCA assay, respectively. Statistical significance between the two groups was determined using an unpaired Student's t-test. **P<0.01, ***p < 0.001. **(C)** Gel agarose electrophoresis to confirm the integrity of the RNA extracted from different samples submitted for mRNA-seq analysis. Three biological replicates (1-3) of control (c) and doxorubicin (dx)-treated cells were included. The molecular size was determined by 1 kb DNA ladder (n3232, New England Biolabs, USA).

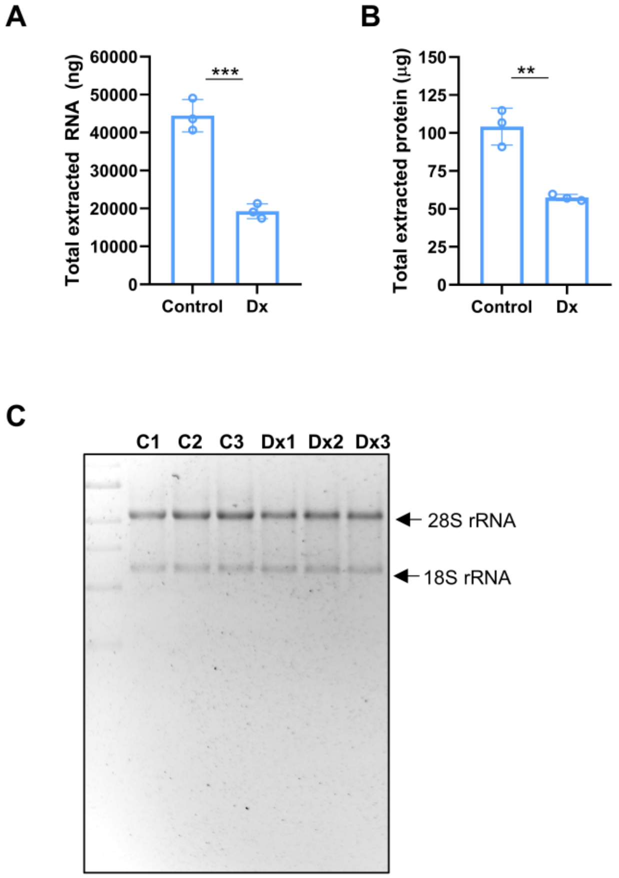
